# Uptake of CD36 and its natural and therapeutic ligands by clathrin-mediated endocytosis

**DOI:** 10.64898/2026.04.06.716751

**Authors:** Hao-Yang Liu, Zhengyu Wang, Rahul Sharma, Jessica Perez, Brandon W. Kusaj, Hongfei Zhou, Minmin Wang, Jon M. Huibregtse, Hong-Yu Li, Jeanne C. Stachowiak

## Abstract

The scavenger receptor CD36 mediates cellular uptake of diverse extracellular ligands including long-chain fatty acids. Recent work has demonstrated that CD36 is also required for uptake of bulky, amphipathic therapeutic compounds including proteolysis-targeting chimeras (PROTACs). However, because the mechanism by which cells internalize CD36 itself remains unknown, how its diverse ligands enter cells has remained mysterious. CD36 has been thought to associate with caveolae, flask-like invaginations of the plasma membrane, yet recent reports have cast doubt on the role of these structures in protein trafficking. Using live cell imaging, we show that CD36 strongly colocalizes with sites of clathrin-mediated endocytosis, where it concentrates both long-chain fatty acids and PROTACs. In contrast, neither CD36 nor its ligands colocalized substantially with caveolae. Further, we identify previously unrecognized binding interactions between CD36 and the clathrin machinery, which help to explain recruitment of CD36 and its ligands to endocytic sites. Importantly, disruption of clathrin assembly inhibits endocytic uptake of both long-chain fatty acids and PROTACs, and abolished PROTAC-induced degradation of multiple target proteins that utilize diverse E3 enzymes. These results suggest that clathrin-mediated endocytosis is essential for internalization of CD36 and its ligands, offering new insights into diverse diseases that rely on CD36 for ligand import, and potentially inspiring new strategies for optimizing targeted protein degraders.

## Introduction

CD36 is a multi-ligand scavenging receptor at the plasma membrane of diverse cell types, with roles in disease pathogenesis and the delivery of novel therapeutics^1–3^. With regard to cardiovascular disease, dysregulated expression of CD36 mediates excessive lipid accumulation, leading to lipotoxicity^4,5^, abnormal insulin response^6,7^, and apoptosis^8^. Meanwhile, in cancerous cells, CD36 facilitates tumor progression by altering lipid metabolism and signaling pathways, including AKT/GSK3β/β-catenin, Wnt, and TGF-β, to promote proliferation, metastasis, and drug resistance^9,10^.

The physiological and clinical importance of CD36 derives from its diverse ligands. Among its endogenous ligands, long-chain fatty acids are the basic units of larger lipid molecules, playing important roles in energy storage, cell membrane structure and signaling^11–13^. CD36 can coordinate fatty acids utilization and link fatty acid-derived signaling molecules to broader metabolic regulation. Beyond its physiological ligands, CD36 was recently found to promote the cellular uptake of proteolysis-targeting chimeras (PROTACs)^14^. PROTACs are a class of engineered bifunctional small molecules that drive selective degradation of pathogenic target proteins through recruitment of ubiquitin E3 ligases^15–17^. PROTACs have demonstrated the potential to eliminate previously “undruggable” targets in prostate and breast cancer, among others, leading to a new and growing class of biopharmaceutic that have attracted substantial investment and are increasingly successful in clinical applications^14,16,18^.

Despite the physiological and therapeutic importance of its ligands, how CD36 facilitates their cellular entry is not well understood. Previous studies have implicated caveolae, flask-shaped invaginations of the plasma membrane, in CD36 internalization^19–21^. However, the contribution of caveolae to endocytosis has been increasingly debated over the past decade. Live-cell imaging data suggest that caveolae are relatively stable on the plasma membrane, implying that endocytic activity may be limited under basal conditions^22,23^. These observations have raised questions about the role of caveolae in constitutive endocytic transport. Similarly, quantitative imaging suggests that a negligible fraction of endosomes are marked by caveolin, in contrast to an overwhelming majority that associated with clathrin^24^. Along the same lines, analysis of endogenously tagged caveolar proteins demonstrated that, while caveolae can depart from the plasma membrane, they contain relatively low levels of bulk plasma membrane proteins, suggesting a specialized role in membrane lipid transport^25^. More specifically, a recent screen for CD36 interacting proteins identified clathrin and the adaptor protein complex AP2, rather than components of caveolae, deepening the uncertainty surrounding the internalization mechanism of CD36^14^. Taken together, these findings motivate a more direct examination of the machinery responsible for CD36 internalization and ligand uptake.

Resolving this question is particularly important for understanding the cellular entry of CD36-dependent ligands, including long-chain fatty acids and amphipathic drug compounds, such as PROTACs. Due to their lipophilicity, fatty acids were initially thought to enter cells via passive diffusion into cellular membranes, without the involvement of protein receptors^26,27^. However, passive uptake appears insufficient to account for the rapid, saturable, substrate-selective, and hormone-regulated nature of long-chain fatty acids uptake observed in many tissues and cell types^28–30^. Although both passive and protein-mediated uptake are currently widely recognized, growing evidence suggested that a substantial portion of fatty acid uptake is mediated by active cellular processes in tissues with high demand^31,32^. PROTACs present a related but distinct challenge, as their large size and physicochemical properties often limit passive membrane permeability, raising questions about how they achieve sufficient intracellular exposure to induce target degradation^33–35^. Although CD36 has been recently shown to facilitate PROTAC uptake^14^, the endocytic machinery underlying this process remains unknown. Therefore, determining the specific mechanism by which CD36 is internalized is necessary to clarify how cells take up both essential nutrients and clinically-important therapeutic molecules.

Here we investigated the mechanism by which CD36 and its natural and therapeutic ligands enter cells. Live cell imaging revealed that CD36 strongly associates with sites of clathrin-mediated endocytosis at the plasma membrane. In contrast, we observed very little association with caveolae. Similarly, long-chain fatty acids and PROTACs associated with clathrin-coated pits in a CD36 dependent manner. We show that these interactions arise from previously unidentified binding motifs within the intracellular domain of CD36 for clathrin adaptor proteins. Strikingly, inhibition of clathrin assembly resulted in reduced cellular uptake of both long-chain fatty acids and PROTACs, and a near complete loss of PROTAC-mediated target degradation across multiple E3 ligases and target proteins. Together these results suggest that CD36 is primarily internalized by clathrin-mediated endocytosis. This finding has broad significance for our understanding of diverse pathologies, from cardiovascular disease and diabetes to cancer, in which CD36 plays an important role in uptake and storage of long-chain fatty acids. Further, we anticipate that knowledge of the CD36 uptake mechanism can be used to optimize and expand upon the design of targeted degraders and other polar therapeutic compounds recently discovered to rely on CD36 for entry into target cells.

## Results

### CD36 associates with clathrin-coated pits at the cellular plasma membrane

To gain insight into the internalization mechanism of CD36, we began by comparing the association between CD36 and markers of the caveolae and clathrin-mediated endocytic pathways during high resolution live cell imaging experiments (Figure 1A). We selected SUM159 human breast cancer-derived epithelial cells because their well spread morphology and thin lamellipodia structures are ideal for visualizing endocytic structures on the plasma membrane. The C terminal cytosolic tail domain of CD36 (residues 462-472) is thought to play an important role in endocytosis^1,19^. Therefore, we fused RFP to the N terminus of CD36 (RFP-CD36) to visualize its spatial distribution on the surface of the plasma membrane, while leaving the C-terminus free to interact with the endocytic machinery. To visualize caveolae, we transiently expressed caveolin1, a major structural component of caveolae (with its C terminus fused to GFP, caveolin1-GFP)^36,37^, and looked for potential colocalization with transiently expressed RFP-CD36. Confocal fluorescence images of the adherent plasma membrane showed that caveolin1-GFP exhibited a discrete punctate distribution on the cell membrane, consistent with the appearance of caveolae in previous studies^37^. RFP-CD36 was likewise enriched in distinct puncta; however, the two sets of puncta had little obvious overlap (Figure 1B), implying that CD36 was not enriched in caveolae. As a positive control for colocalization with caveolin1-GFP, we used cavin1, a second structural component of caveolae^36^. As expected, RFP tagged cavin1 (cavin1-RFP) formed punctate structures on the plasma membrane that strongly colocalized with caveolin1-GFP (Figure 1C).

**Figure 1.**
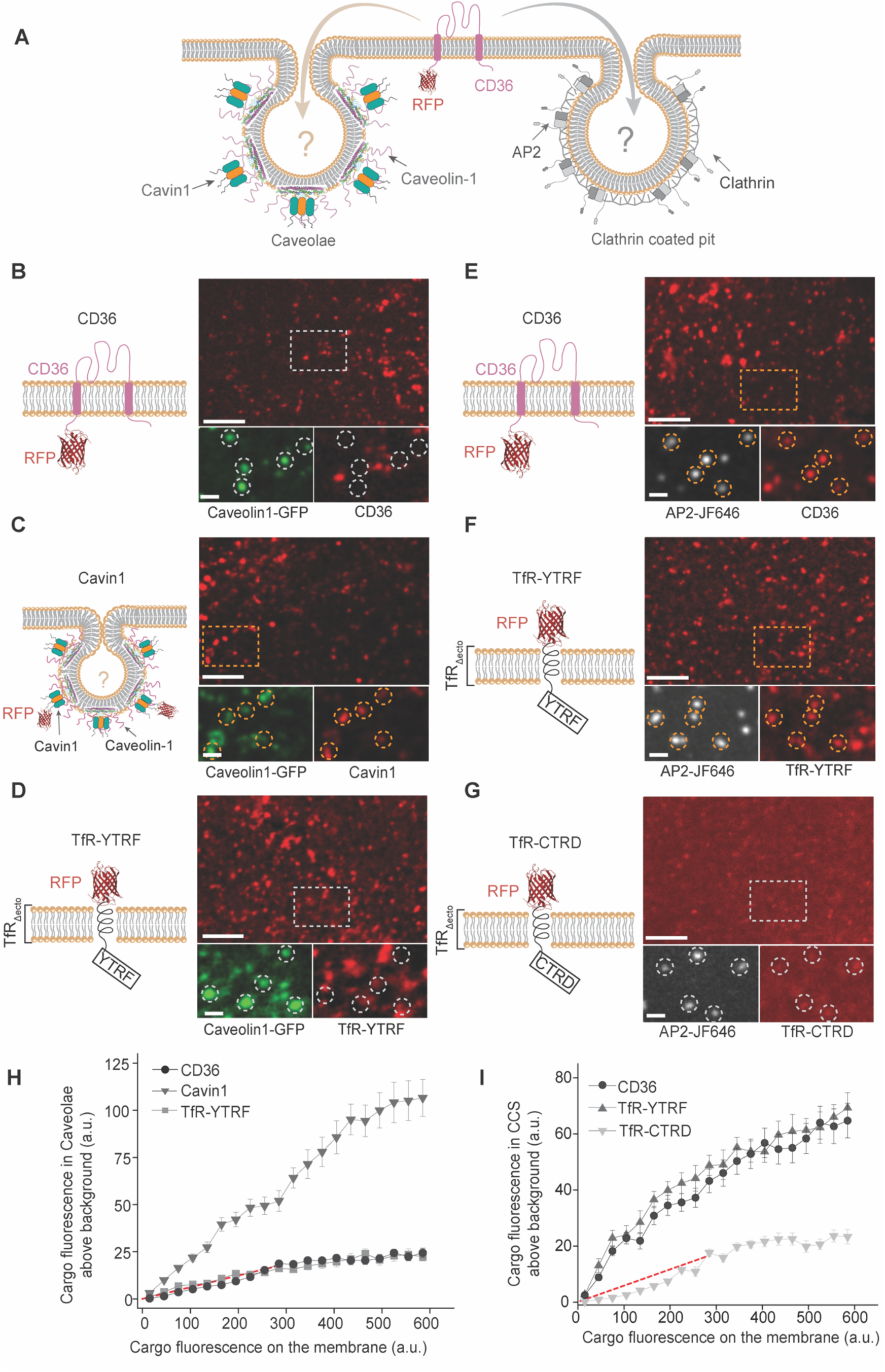
CD36 selectively partitions into clathrin coated pits rather than caveolae. (A) Schematic illustration depicting the two hypothesized endocytic routes for CD36 internalization: clathrin-mediated endocytosis (left, involving AP2 and clathrin) versus caveolin-dependent endocytosis (right, involving caveolin1 and cavin1). (B-D) Spinning disk confocal images of the plasma membrane of SUM 159 cells transiently expressing caveolin1-GFP (green) along with RFP-CD36 (B), the caveolae specific cargo Cavin1-RFP or clathrin specific cargo TfR-YTRF. (E–G) Spinning disk confocal images of the plasma membrane of SUM 159 cells labeled with AP2-JF646 along with the expression of CD36 (E), the clathrin specific cargo TfR-YTRF (F) or the clathrin-independent negative control TfR-CTRD (G). Dashed boxes indicate regions magnified in the lower insets. Yellow and white dashed circles denote endocytic structures with and without colocalized signals, respectively. (H) The relative number of transmembrane fusion proteins localized in caveolae structures is shown against the relative concentration of fusion proteins on the plasma membrane around each structure. Data were collected from over 5000 endocytic sites in n=20 independent cells expressing the CD36-RFP, cavin1-GFP or TfR-YTRF-RFP model protein. Each point reflects the average value from caveolae structures binned by the local membrane concentration of the proteins. (I) The relative number of transmembrane fusion proteins localized in clathrin coated structures is shown against the relative concentration of fusion proteins on the plasma membrane around each pit. Data were collected from over 6000 endocytic sites in n=25 independent cells expressing the CD36, TfR-YTRF or TfR-CTRD model protein. Each point reflects the average value from clathrin coated structures binned by the local membrane concentration of the proteins. The red dashed lines in panel H and I indicate the level of correlation expected from incidental colocalization, as determined from Figure S1. Scale bars: 5 µm (main images) and 500 nm (insets). Error bars denote mean ± s.e.m.

The limited colocalization of CD36 with caveolin-1 prompted us to examine whether CD36 instead associates with sites of clathrin-mediated endocytosis (CME), the major endocytic carriers for specific uptake of proteins from the plasma membrane^38^. To track sites of CME during live cell imaging, we employed a SUM159 cell line in which genome editing was used to fuse a HaloTag at both alleles of the *μ*2 subunit within the AP2 complex, the major adaptor protein complex of the clathrin pathway^39^. We chose AP2 rather than clathrin itself because AP2 is restricted to the plasma membrane, whereas clathrin also decorates internal organelles such as endosomes and the Golgi^40^. Given the known 1:1 stoichiometry between AP2 and clathrin, tracking AP2 provides high confidence for identification of clathrin coated structures at the plasma membrane^40^. For visualization of AP2 during fluorescence imaging, we used the far red HaloTag ligand JF646^41^. Live-cell imaging revealed clear colocalization between RFP-CD36 puncta and puncta enriched in AP2 at the plasma membrane, implying robust recruitment of CD36 to sites of clathrin-mediated endocytosis (Figure 1E). As a positive control for recruitment to clathrin coated pits, we used TFR-YTRF, a model receptor consisting of the transmembrane and cytosolic domains of the transferrin receptor (TFR), fused to an extracellular RFP. As the YTRF motif in the cytoplasmic domain of this model receptor can directly bind to the μ2 subunit of AP2, it is internalized almost exclusively via CME^42,43^. As expected, TfR-YTRF and AP2 exhibited strong colocalization (Figure 1F). Similar to CD36, TfR-YTRF showed almost no colocalization with caveolin1-GFP, providing a complementary negative control for caveolae localization (Figure 1D). To assess the specificity of its recruitment to AP2 positive sites, we also examined the TfR-YTRF model receptor mutant, TfR-CTRD, which has previously been shown to exhibit significantly reduced localization to sites of clathrin-mediated endocytosis^44–46^. As expected, TfR-CTRD showed much weaker enrichment in AP2 positive structures, in comparison to CD36 and TfR-YTRF (Figure 1G).

To quantify the differential partitioning of CD36 and the control markers (cavin1, TfR-YTRF, TfR-CTRD) to caveolae versus sites of CME, we used a publicly available algorithm that quantifies colocalization among punctate structures in fluorescence images^47^. The algorithm first identifies individual punctate structures by applying a 2D Gaussian fit to puncta in the master channel (caveolin1 or AP2). The algorithm then fits the corresponding signal in the subordinate channel (CD36, cavin1, TfR-YTRF, TfR-CTRD) to determine enrichment relative to the local background. The resulting Gaussian amplitude reflects the relative abundance of the subordinate protein within each clathrin coated pit or caveolae site^48^. At the same time, the software measures the average fluorescence intensity of the surrounding area to gauge the local concentration of the subordinate protein. Using these data, we plotted the relative recruitment of each protein into individual endocytic structures (caveolae or clathrin coated pits) as a function of its local relative concentration. Before presenting these data, it is important to note that, when comparing any two patterns of closely-spaced puncta, there will be some rate of non-specific colocalization. This effect was evident when we used AP2 as the master channel and caveolin1 as the subordinate channel (Figure S1A) or vice versa (Figure S1B), which yielded a modestly increasing trend with a slope slightly above six percent (Figure S1A). Given that these proteins are known to be components of entirely orthogonal endocytic pathways with no overlap in protein machinery, the slope of the resulting trends approximate the extent of non-specific colocalization, a threshold that we would expect true colocalization events to substantially exceed. Having established this threshold, we applied the same approach to examine relative enrichment of cavin1, CD36, TfR-YTRF, and TfR-CTRD in caveolae and sites of CME.

As expected, the resulting plots showed strong enrichment of cavin1 within caveolae as a function of local cavin1 concentration, with a slope that substantially exceeded the non-specific threshold (red dotted line). In contrast, TFR-YTRF exhibited significantly lower enrichment levels across the measured range, with a slope similar to the non-specific threshold. As noted above, this result is expected because TFR-YTRF is known to be strongly and exclusively internalized by CME. Similarly, CD36 exhibited only non-specific colocalization with caveolin1 (Figure 1H). In contrast, at AP2 positive endocytic sites, CD36 exhibited a strong trend of increasing colocalization with increasing local concentration, well above the non-specific threshold. The resulting trend had a similar slope to that of TFR-YTRF, the positive control for uptake by CME. In contrast, TFR-CTRD, the negative control, had a much lower slope, consistent with the non-specific threshold, as expected (Figure 1I). These results suggest that CD36 associates preferentially with CME sites over caveolae to a degree that is similar to the transferrin receptor, one of the best-established and most specific cargo proteins of the clathrin pathway.

### Molecular mechanism of CD36 recruitment to clathrin-coated structures

The observation that CD36 concentrates at sites of CME motivated us to investigate the molecular mechanisms underlying the interaction between CD36 and the protein machinery responsible for clathrin-mediated endocytosis. The C-terminal tail of CD36 (residues 462-472) is known to undergo multiple post-translational modifications and contains potential sorting signals that may interact with the endocytic machinery^49–51^. To systematically dissect the contribution of these motifs, we constructed a series of mCherry tagged variants of CD36 (mCherry-CD36) with mutations or truncations in the C terminal domain and expressed them in the SUM159 cell line used above (Figure 2A-D). To quantify their recruitment, we plotted the relative concentration (intensity) of each CD36 variant per endocytic structure against its local concentration (intensity) on the cell membrane (Figure 2E).

**Figure 2.**
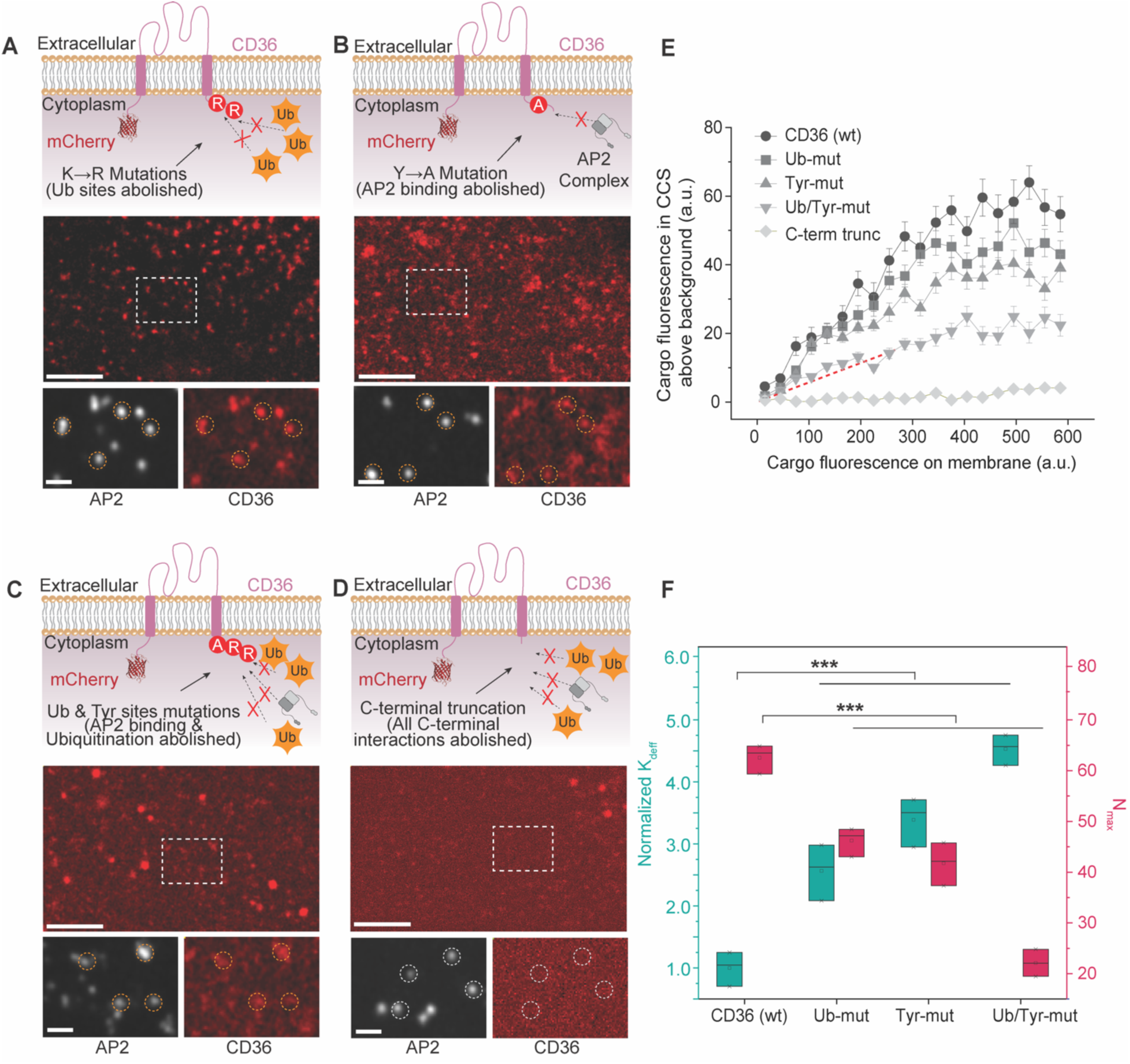
Molecular mechanism of CD36 recruitment to clathrin-coated structures. (**A–D**) Schematics and representative spinning disk confocal images of SUM 159 cells expressing mCherry-tagged CD36 variants with specific mutations in the cytosolic tail: (**A**) the ubiquitination-deficient mutant (Ub-mut), where lysine residues (K) are mutated to arginine residues (R); (**B**) the AP2-binding-deficient mutant (Tyr-mut), where tyrosine residues (Y) are mutated to alanine residues (A); (**C**) the double mutant (Ub/Tyr-mut) combining both modifications; and (**D**) the C-terminal truncation mutant (C-term trunc) lacking the entire C-terminal cytosolic domain. Top diagrams illustrate the specific molecular disruptions. Bottom images show the plasma membrane, where white circles highlight sites of clathrin-mediated endocytosis marked by AP2-JF646 (white), and red fluorescence indicates the distribution of the indicated CD36 variants. Scale bars are 5 µm (main image) and 2 µm (insets). (**E**) The relative number of transmembrane model proteins localized in clathrin-coated structures is shown against the relative concentration of model proteins on the plasma membrane around each structure. Data were collected from 6552 endocytic sites in 25 cells expressing the CD36 model protein, 7147 endocytic sites in 25 cells expressing the Ub-mut model protein, 6428 endocytic sites in 22 cells expressing the Tyr-mut model protein, 8061 endocytic sites in 24 cells expressing the Ub/Tyr-mut model protein, and 7089 endocytic sites in 20 cells expressing the C-term trunc model protein. Each point reflects the average value from clathrin-coated structures binned by the local membrane concentration of the proteins. The red dashed lines indicate the level of correlation expected from incidental colocalization, as determined from Figure S1. Error bars denote mean ± s.e.m. (**F**) Box plot showing the relative K_deff_ values (green, left y axis) and N_max_ values (red, right y axis) for the CD36 (wt), Ub-mut,Tyr-mut and Ub/Tyr-mut proteins. K_deff_ represents the effective dissociation constant, normalized to the CD36 (wt), while N_max_ represents the maximum local concentration of fusion proteins in clathrin-coated structures. Each box displays the median, interquartile range. Data represent mean ± s.e.m.. ***P < 0.001.

We first examined the role of ubiquitylation, as ubiquitin binding adaptor proteins during CME can specifically recognize and concentrate ubiquitylated cargo at endocytic sites^52,53^. Lysine residues K469 and K472 in the C terminal domain of CD36 have previously been identified as ubiquitylation sites^54^. We therefore mutated these two residues to arginine (Ub-mut, K469R/K472R, Figure 2A). The resulting plots revealed a significant reduction of approximately 24% in the relative enrichment of the resulting mutant (Ub-mut) at sites of CME compared to wild type CD36 (Figure 2E). But the majority of recruitment remained, implying that other interactions in addition to ubiquitination contribute to CD36 localization at endocytic sites.

To further investigate these additional interactions, we examined the potential influence of direct adaptor recognition. Tyrosine based sorting signals typically follow the YXXΦ motif, where X represents any amino acid and *Φ* represents a bulky hydrophobic residue, and are recognized by the *μ*2 subunit of AP2^55^. The YTRF motif in the transferrin receptor is a well characterized example^55,56^. While CD36 lacks a canonical YXXΦ motif, its C-terminal tail contains a YCAC sequence beginning at Y463, which we hypothesized might function as a non-canonical tyrosine based sorting signal. To test this possibility, we mutated Y463 to alanine (Tyr-mut, Y463A, Figure 2B). Images of localization to endocytic sites suggest that this mutant exhibited a significant reduction in enrichment at sites of clathrin-mediated endocytosis compared to Ub-mut (Figure 2B), consistent with the quantification in Figure 2E.

To determine if these two mechanisms operate independently or additively, we characterized a triple mutant combining both types of mutations (Ub/Tyr-mut, Y463A/K469R/K472R, Figure 2C). When compared to each of the previous mutants, the puncta intensity of the triple mutant (Ub/Tyr-mut) in the endocytic structure appears to be further diminished, implying that these two mutations have an additive effect. Consistent with these images, the quantification showed that the combination of the K to R mutations and the Y to A mutation yielded a low level of CD36 enrichment at CME sites that was comparable to the non-specific threshold (red dotted line), suggesting that these two types of interactions are sufficient to explain the majority of CD36 localization to CME sites, Figure 2C. Further removal of the entire C terminal domain of CD36 resulted in a diffuse distribution at the plasma membrane, largely devoid of puncta (Figure 2D), such that the slope of the resulting curve fell well below the non-specific threshold (Figure 2E). This further decrease suggests that additional weak binding interactions encoded within the C-terminal domain may contribute to endocytosis or clustering of CD36 by unknown mechanisms.

In Figure 2E, recruitment of each of the CD36 mutants rose with increasing plasma membrane concentration, approaching a plateau value that represents the saturated capacity of CME sites for each mutant^48^. To quantify the resulting trends, we applied a simplified multivalent binding model (Equation 1)^48^. In this model, the mean occupancy of each endocytic structure by transmembrane proteins, represented by ⟨*n*⟩, is a function of the maximum capacity per structure (N_max_), the relative concentration of the transmembrane proteins on the plasma membrane (C_mem_), and the effective dissociation constant for transmembrane protein binding to endocytic sites (K_deff_), Equation 1.

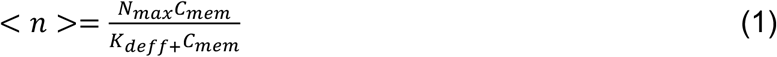

We fit the data in Figure 2E with this equation, leaving K_deff_ and N_max_ as free parameters (Figure 2F). This analysis revealed that, compared to wild-type CD36 (WT), the effective dissociation constant K_deff_ of each variant exhibited a stepwise increase, indicating weakening binding affinity for CME sites. Specifically, the ubiquitinated mutant (Ub-mut) increased by 2.6-fold, the tyrosine mutant (Tyr-mut) increased by 3.4-fold, and the dual mutant Tyr/Ub-mut increased by 4.5-fold compared to wild-type CD36. Interestingly, we also observed a concurrent substantial reduction in saturation capacity N_max_. These results suggest that weaker binding of the CD36 mutants to the endocytic machinery allowed other endogenous receptors to compete with them for limited space within CME sites. Together these findings suggest that CD36 relies on multivalent interactions to achieve high-affinity recruitment to clathrin-coated structures. The reliance of CD36 on multiple weak interactions, rather than stronger canonical binding motifs^38^, helps to explain why clathrin-mediated uptake of CD36 was not described previously. Having identified key residues in the C-terminal domain of CD36 that mediate localization to CME sites, we next sought to evaluate the dependence of ligand uptake on CD36 and clathrin-mediated endocytosis.

### Long chain fatty acids colocalize with CD36 at sites of clathrin-mediated endocytosis

Next, we investigated whether long-chain fatty acids, among the major types of natural ligands of CD36, are recruited to sites of clathrin-mediated endocytosis (Figure 3A). To visualize their distribution at the plasma membrane, we synthesized a fluorescent stearic acid analogue bearing ATTO488 at the ω end of the hydrocarbon chain through an amide linkage, leaving the carboxyl group unmodified (Figure 3B; Scheme S1 and Figures S5, S12, S14 and S16). We optimized the design to place the fluorophore as far as possible from the carboxyl group thought to be involved in CD36 recognition^57,58^, thereby minimizing interference with receptor binding. At the same time, we utilized the hydrophilic dye ATTO488 to minimize dye-driven membrane partitioning of the probe^59^. First, we examined cells exposed to fluorescently labeled CD36 after adding ATTO488-labeled stearic acid from a DMSO stock solution. Distinct fluorescent puncta in the ATTO 488-stearic acid channel were observed on the cell surface and showed significant colocalization with CD36 and sites of CME (AP2) (Figure 3C). Next, we simultaneous visualized caveolae (caveolin1-mcherry) and sites of CME (AP2-JF646). After addition to these cells, stearic acid partitioned clearly into sites of CME and was largely absent from caveolae (Figure 3D).

**Figure 3.**
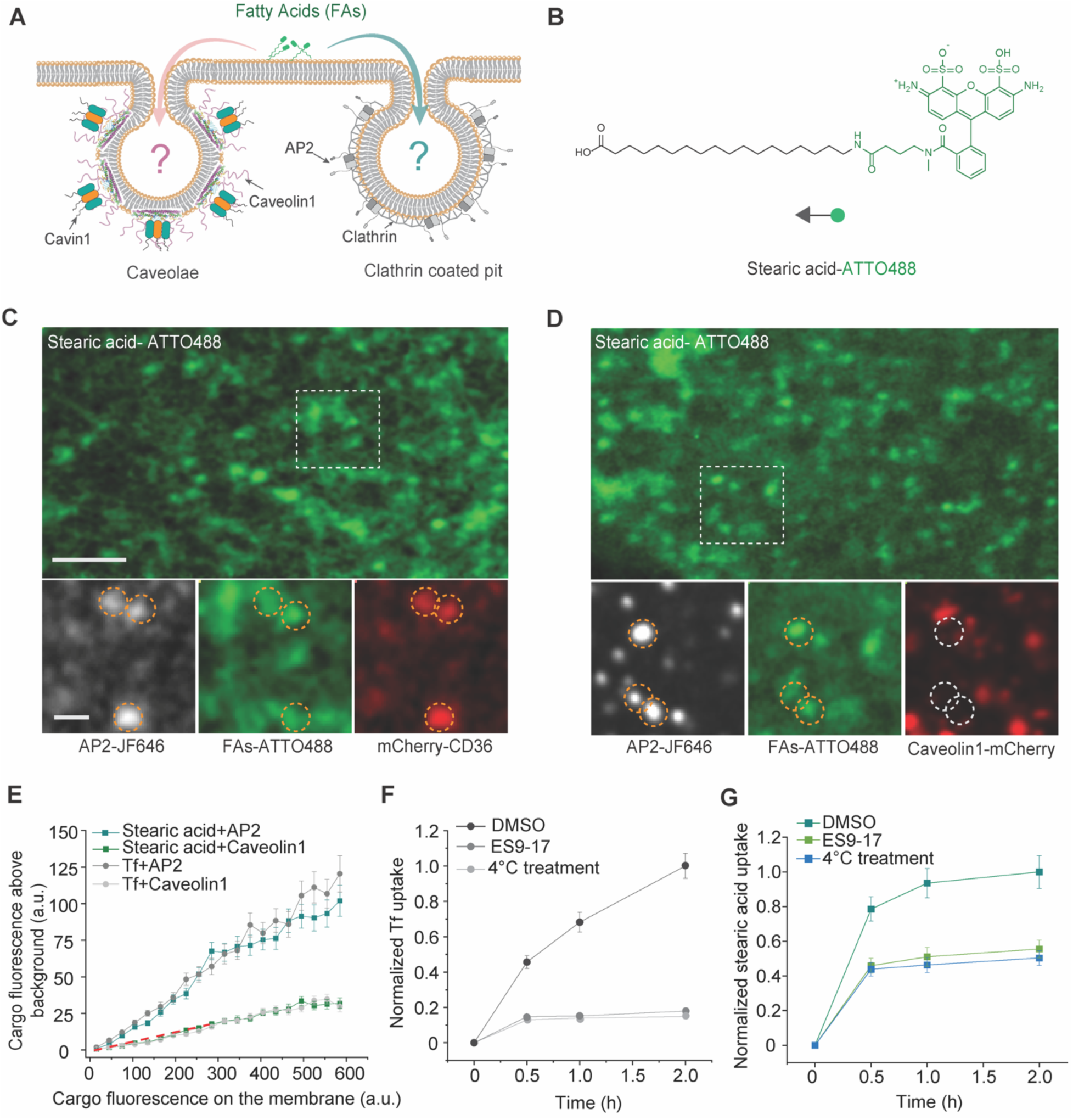
Long chain fatty acids colocalize with CD36 at sites of clathrin-mediated endocytosis. (A) Schematic illustrating the potential involvement of caveolae and clathrin-coated pits in long-chain fatty acid uptake. Caveolin1 and Cavin1 mark caveolae, whereas AP2 and clathrin are components of clathrin-coated pits. Arrows and question marks indicate the candidate uptake routes investigated in this study. (B) Chemical structure of the fluorescently labeled long-chain fatty acid, Stearic acid-ATTO488, serving as a physiological model ligand for CD36. (C) Representative spinning disk confocal images of SUM 159 cells expressing mCherry-CD36 (red) and treated with Stearic acid-ATTO488 (green). Clathrin-coated structures (CCS) are marked by AP2-JF646 (white). Orange dashed circles highlight the robust triple colocalization of CD36, fatty acids, and AP2 at endocytic sites, indicating efficient cargo recruitment to the clathrin machinery. Scale bars are 5 µm (main images) and 500 nm (insets). (D) Confocal images of cells expressing a caveolar marker (e.g., Caveolin1-mCherry, red) treated with Stearic acid-ATTO488 (green) and labeled with AP2-JF646 (white). Orange dashed circles indicate the partitioning of fatty acids into AP2-positive sites, while white dashed circles highlight the distinct, non-overlapping caveolar structures. This demonstrates that fatty acids are actively excluded from caveolae. (E) Quantification of stearic acid-ATTO488 and fluorescently labeled transferrin (Tf) enrichment at CME sites and caveolae. Cargo fluorescence above local background is plotted against cargo fluorescence intensity in the surrounding plasma membrane. Transferrin served as a positive control for localization to clathrin-coated structures and a negative control for caveolar localization. The red dashed lines indicate the level expected from incidental colocalization, as determined from Figure S1. Data were collected from over 5000 endocytic sites in n=20 independent cells respectively. Data represent mean ± SEM. (F) Time course of Tf-ATTO488 uptake in DMSO (control) and 10μM ES9-17-treated SUM159 cells, measured by flow cytometry at the indicated time points. Tf-ATTO488 uptake was also measured at 4 °C to broadly inhibit active processes, including endocytosis. (G) Time course of Stearic acid-ATTO488 uptake in DMSO (control) and 10μM ES9-17-treated SUM159 cells, measured by flow cytometry at the indicated time points. Stearic acid-ATTO488 uptake was also measured at 4 °C to broadly inhibit all forms of endocytosis.

Using the image analysis procedure described above, we quantified stearic acid fluorescence above local background at sites of CME and caveolae as a function of stearic acid fluorescence intensity in the surrounding plasma membrane. Transferrin (Tf), labeled with ATTO 488 was used as a positive control for internalization at sites of CME and as a negative control for internalization by caveolae^60^ (Figure S2). As expected, the resulting plots showed that transferrin exhibits significant enrichment in AP2-positive structures, whereas at caveolin1 positive sites, its signal does not exceed the threshold for incidental colocalization (Figure 3E, red dotted line). Stearic acid-ATTO488 exhibited a similar pattern, with strong recruitment to sites of CME and only incidental recruitment to caveolae (Figure 3E). These results suggest that stearic acid has a bias toward internalization through CME, rather than caveolae, which is just as strong as that of transferrin, a ligand that is thought to rely almost exclusively on the clathrin pathway.

To assess uptake and accumulation of the ligands in cells, we used flow cytometry to compare the increase in ATTO 488-labeled ligand intensity over time for both transferrin and stearic acid. For transferrin, we observed an increase in intensity that continued to rise over the two-hour measurement period. Pretreatment of cells with ES9-17, a drug that blocks CME by specifically inhibiting assembly of the clathrin lattice^61^, resulted in a dramatic loss of transferrin fluorescence, approximately 82% at the two hour timepoint (Figure 3F). A similar loss of transferrin uptake was observed when cells were treated at 4°C, which is known to severely attenuate clairin-mediated endocytosis and most other active cellular processes^62^. The stearic acid intensity rose more abruptly, approaching its plateau value within an hour. Treatment with ES9-17 resulted in reduced stearic acid signal at each time point, but the reduction was below 50% (46% at 2h, Figure 3G). The residual signal could be due to either passive adsorption and incorporation of stearic acid directly onto the plasma membrane, as has been previously observed^63^ or uptake through a clathrin-independent endocytic pathways. To assess the influence of passive incorporation, we exposed cells to stearic acid at 4 °C, where the resulting signal decreased to a level very similar to that resulting from ES9-17 treatment at 37°C, suggesting that the major active process involved in stearic acid uptake is CME. Collectively, these results indicate that CME, rather that caveolae, is responsible for endocytic uptake of stearic acid.

### CD36 is essential for the recruitment of PROTACs to sites of clathrin-mediated endocytosis

We next examined the endocytic mechanisms responsible for uptake of PROTACs, a growing class of therapeutic compounds, which have recently been shown to rely on binding interactions with CD36 for internalization by cells. We began by investigating the colocalization of fluorescently labeled PROTACs with CD36 and endocytic sites on the plasma membrane. For this purpose, we synthesized a fluorescently labeled PROTAC molecule, BRD4-VHL-ATTO488 (Scheme S2 and Figures S6–S11, S13, S15, and S17), which targets the transcriptional regulator, bromodomain-containing protein 4 (BRD4) for degradation by linking it to the ubiquitin E3 ligase, von Hippel Lindau (VHL) (Figure 4A)^64,65^. Multi-channel live-cell imaging demonstrated a clear colocalization between the PROTAC (ATTO488), CD36 (mCherry) and sites of CME (AP2-JF646) (Figure 4B). In contrast, BRD4-VHL-ATTO488 showed minimal colocalization with caveolae (Cav1-mCherry), which did not exceed the level of incidental colocalization (Figure S3).

**Figure 4.**
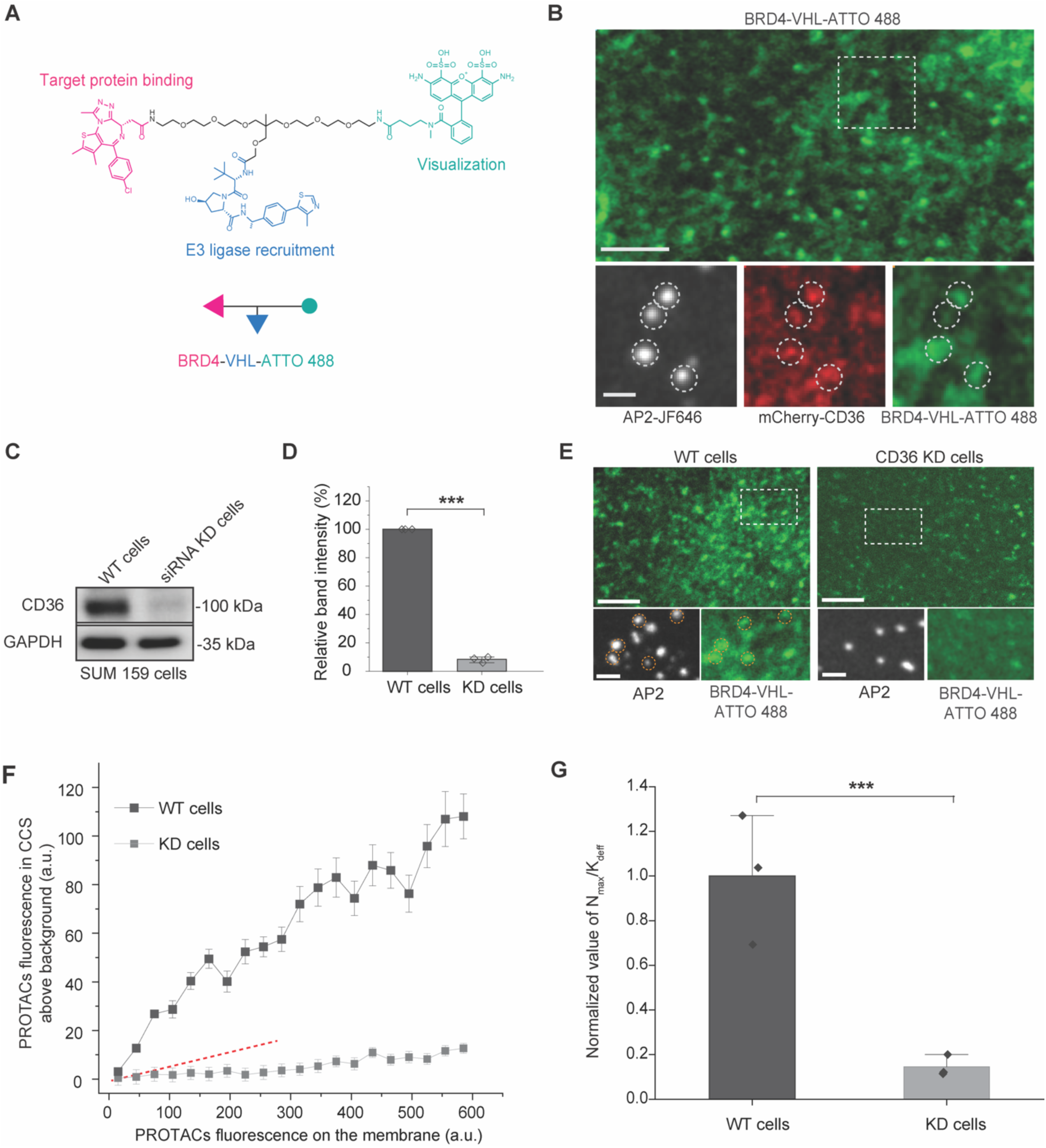
CD36 is essential for the recruitment of PROTACs to sites of clathrin-mediated endocytosis. (**A**) Chemical structure of the fluorescently labeled PROTAC, BRD4-VHL-ATTO 488. (**B**) Representative spinning disk confocal images showing the recruitment of PROTACs to CD36-positive endocytic sites. SUM 159 cells expressing CD36 model receptors were treated with BRD4-VHL-ATTO 488. White dashed circles highlight the triple colocalization of AP2-JF646 (white), CD36 model receptor (red), and the PROTACs (green). Scale bars are 5 µm (main image) and 1 µm (insets). (**C**) Western blot analysis of whole-cell lysates from SUM 159 cells treated with non-targeting control siRNA (WT cells) or CD36-targeting siRNA (KD cells). The blot confirms the specific depletion of CD36, with GAPDH serving as a loading control. (**D**) Quantification of CD36 band intensity from (A), normalized to GAPDH levels. (**E**) Representative spinning disk confocal images of the plasma membrane of WT and CD36 KD cells treated with BRD4-VHL-ATTO 488 (green). Sites of clathrin-mediated endocytosis are marked by AP2-JF646 (white). White dashed circles in the magnified insets highlight endocytic sites. Note the accumulation of PROTACs at sites of clathrin-mediated endocytosis in WT cells and the absence of such enrichment in CD36 KD cells. Scale bars are 5 µm (main image) and 2 µm (insets). (**F**) Quantification of PROTAC recruitment at sites of clathrin-mediated endocytosis. The fluorescence intensity of the PROTAC within sites of clathrin-mediated endocytosis is plotted against its local membrane fluorescence intensity. Recruitment is robust in WT cells but is abrogated in CD36 KD cells. The red dashed lines indicate the level of correlation expected from incidental colocalization, as determined from Figure S1 (**G**) Normalized recruitment efficiency of PROTACs to sites of clathrin-mediated endocytosis, calculated as the relative ratio of N_max_ to K_deff_ for PROTACs, derived from the curves in (**F**). The data demonstrate that CD36 is essential for the partitioning of PROTACs into the clathrin endocytic pathway. Data represent mean ± s.e.m.. ***P < 0.001.

To determine whether CD36 was required to recruit PROTACs to sites of CME, we knocked down endogenous CD36 in SUM159 cells, as confirmed by western blot (∼90%, Figure 4C, D). CD36 knockdown cells were then incubated with BRD4-VHL-ATTO488. Live cell imaging revealed a significant visual contrast: in wild type cells, PROTACs exhibited significant colocalization with CME sites, which was largely absent in CD36 knock down cells (Figure 4E). Quantitative analysis confirmed that PROTAC recruitment to sites of CME in knock down cells remained at the incidental level (red dotted-line), despite the presence of PROTACs near the plasma membrane. In contrast, images of wild type cells revealed strong recruitment of PROTACs to sites of clathrin-mediated endocytosis (Figure 4F). To quantitatively compare the recruitment efficiency of PROTACs, we calculated the slope of the recruitment curves (N_max_ / K_deff_), which provides a measure of recruitment efficiency that takes into account both the capacity and affinity of endocytic sites for PROTACs. As shown in Figure 4G, the slope was reduced in CD36 KD cells by 86% compared to WT cells, illustrating a strong dependence of PROTAC recruitment on CD36. Collectively, these data indicate that CD36 is required for recruitment of PROTACs to sites of CME.

### Disruption of clathrin-mediated endocytosis blocks cellular uptake of PROTACs

We next examined the accumulation of fluorescent-labeled PROTACs in cell over time using flow cytometry. Using ES9-17 to block CME, we observed a potent reduction in cellular uptake of transferrin (Figure 5B). Similarly, uptake of the fluorescently labeled PROTAC (BRD4-VHL-ATTO488) was reduced by approximately 85% upon exposure to 10 μM ES9-17 for 2h, suggesting a strong dependence on CME. As an additional confirmation, we employed a second strategy to block CME, which was to overexpress a dominant negative protein, the C-terminal fragment of the clathrin adaptor protein, AP180 (AP180ct, residues 516-898). This fragment contains potent clathrin binding domains but lacks membrane binding domains that sequester clathrin in the cytosol, preventing CME^66^. As confirmation of this effect, cells that expressed AP180ct showed a marked reduction in PROTAC fluorescence compared to surrounding cells that did not express the dominant negative protein (Figure 5A). Similar to ES9-17 treatment, expression of AP180ct resulted in a dramatic reduction of cellular uptake for both transferrin and the labeled PROTAC in flow cytometry experiments (Figure 5B). Further, this effect was visualized in live cell imaging experiments at fixed time points (0, 30, 60 minutes, Figure 5C). Together, these data suggest that uptake of the fluorescent-labeled PROTAC is strongly dependent on clathrin-mediated endocytosis.

**Figure 5.**
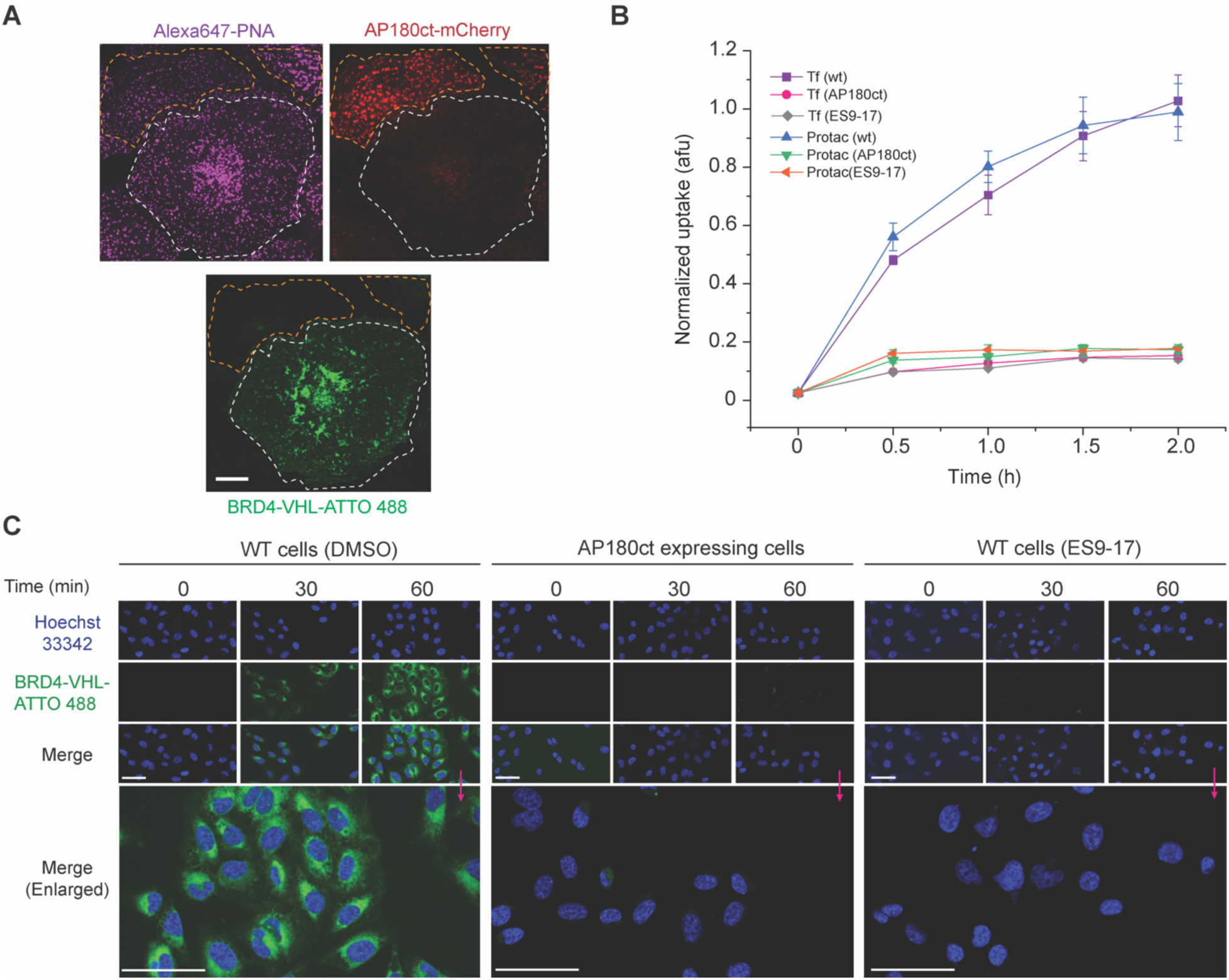
Disruption of clathrin-mediated endocytosis blocks PROTAC uptake. (A) Representative confocal images of SUM-159 cells expressing the CME inhibitor AP180ct-mCherry (red) and incubated with BRD4-VHL-ATTO 488 (green) and Alexa647-PNA (magenta). The expression of AP180ct serves as a dominant-negative perturbation to block the assembly of clathrin. Yellow dashed lines outline AP180ct-expressing cells, which show minimal PROTACs uptake, whereas white dashed lines delineate non-expressing control cells with robust PROTACs signals. Scale bar, 10 μm. (B) Time-course quantification of the internalization kinetics of Tf-ATTO488 (positive control) and BRD4-VHL-ATTO 488 in wild-type (WT) and AP180ct-expressing cells. The uptake of both TfR and the PROTACs is significantly attenuated upon inhibition of the clathrin machinery, afu, arbitrary fluorescence units. Data represent mean ± s.e.m. from n=3 independent experiments. (C) Representative live-cell fluorescence images showing the time-dependent intracellular accumulation of BRD4-VHL-ATTO 488 (green). SUM 159 cells were visualized under three conditions: WT cells treated with DMSO (control), cells expressing the genetic inhibitor AP180ct, and WT cells treated with the chemical clathrin inhibitor ES9-17 (10 μM). Nuclei were counterstained with Hoechst 33342 (blue). Bottom panels show enlarged views of the 60 min time point (indicated by magenta arrows), highlighting robust PROTAC internalization in control cells and lack of accumulation in clathrin-inhibited cells.

### Degradation of target proteins by PROTACS requires clathrin-mediated endocytosis

To evaluate the role of clathrin-mediated endocytosis in target degradation by PROTACs, we used western blots to assess the relative amount of target protein remaining in cells as a function of PROTAC concentration, in the absence and presence of clathrin pathway inhibitors. Specifically, ES9-17 was used to inhibit CME while measuring the impact on target degradation across a panel of different PROTACs. We first evaluated BRD4-VHL-ATTO488, which targets BRD4 by bringing it into contact with the E3 ligase, VHL, as described above. In the DMSO treated control group, BRD4 levels exhibited a robust dose-dependent decrease after 4h treatment, with degradation efficiencies reaching 77% and 91% at 10 nM and 50 nM, respectively (Figure 6A-B). In contrast, pretreatment with 10 μM ES9-17 resulted in BRD4 band intensities for both doses that were comparable to control cells in the absence of BRD4-VHL-ATTO488, indicating that clathrin-mediated endocytosis is required for degradation of BRD4 by this PROTAC. Consistent with these results, the genetic inhibition of CME via AP180ct expression similarly blocked BRD4 degradation (Figure S4), further confirming reliance on the clathrin pathway.

**Figure 6.**
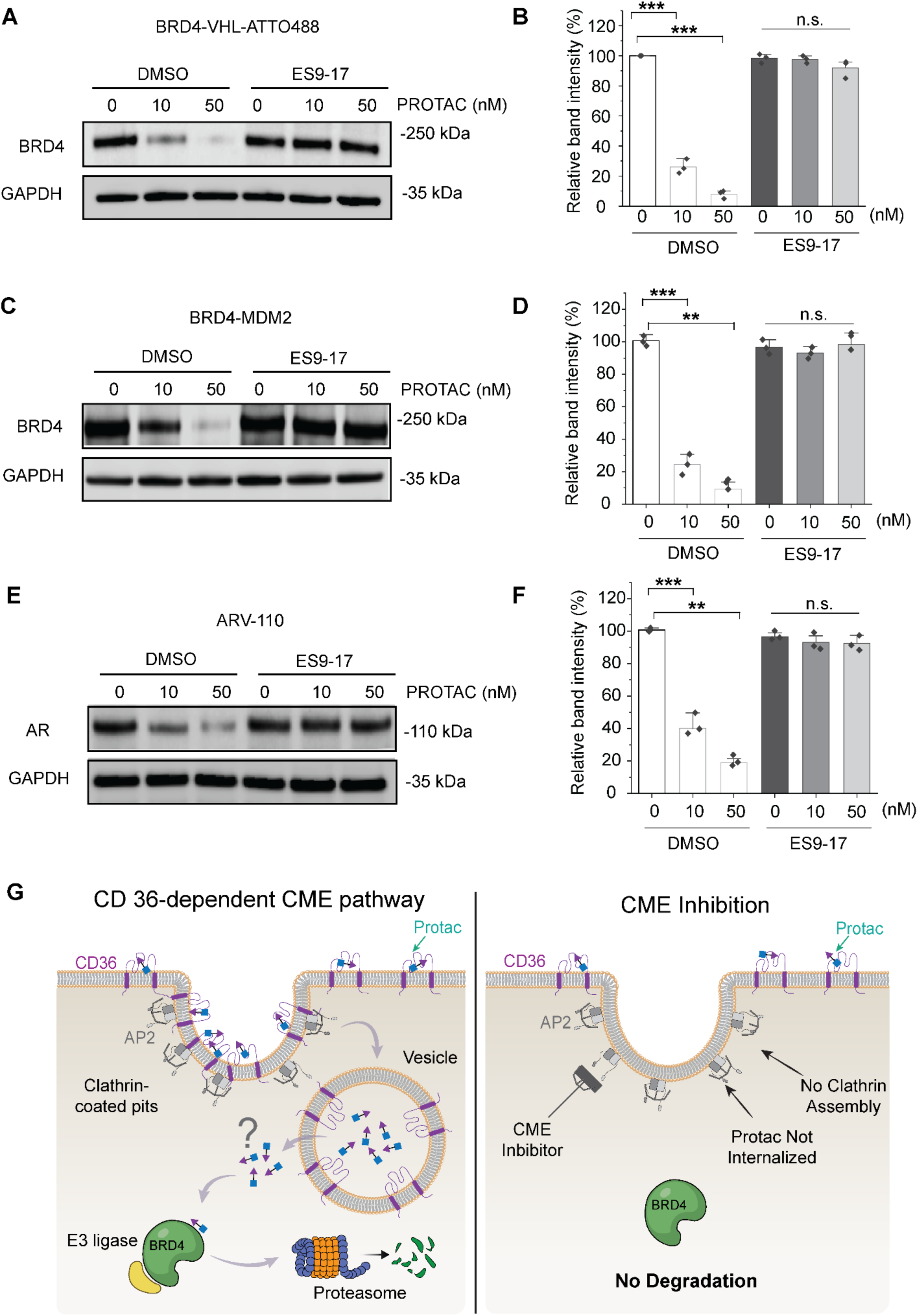
Degradation of target proteins by PROTACS requires clathrin-mediated endocytosis. (**A-F**) Representative Western blots and corresponding quantitative analyses are shown for cells treated with indicated concentrations (0, 10, 50 nM) of PROTACs (**A, B**) BRD4-VHL-ATTO488, (**C, D**) BRD4-MDM2, and (**E, F**) ARV-110 (targeting AR) in the presence of either DMSO or the CME inhibitor ES9-17 (10 μM). Band intensities were normalized to GAPDH. Data are presented from three independent experiments. ** P < 0.01, *** P < 0.001, n.s., not significant. (**G**) Mechanistic model of CD36-dependent PROTAC internalization and activity. (Left) Under physiological conditions, the CD36-PROTAC complex is recruited into clathrin-coated pits, leading to efficient vesicle-mediated cytosolic delivery and subsequent target protein degradation via the E3 ligase-proteasome system. (Right) Chemical inhibition of clathrin assembly (CME Inhibition) traps PROTACs at the cell surface, preventing internalization and thereby abolishing their degradative capability.

To examine whether the requirement for CME depends on the specific E3 ligase recruited, we tested BRD4-MDM2 (A1874), which uses the mouse double minute 2 homolog (MDM2) E3 ligase to degrade BRD4. Similar to BRD4-VHL-ATTO488, degradation of BRD4 by A1874 was blocked upon loss of CME (Figure 6C-D). These results demonstrated that the reliance of PROTACs on CME extends to PROTACs that use different E3 ligases.

Lastly, to determine whether the requirement for clathrin-mediated endocytosis extends beyond BRD4 as a target protein, we investigated the dependence of a third PROTAC, ARV-110 on the clathrin pathway. ARV-110 targets the androgen receptor (AR), a hormone-dependent transcription factor, by recruiting the E3 ubiquitin ligase, Cereblon (CRBN)^67^. Consistent with our results for the other two PROTACs, the ability of ARV-110 to drive degradation of AR was lost when clathrin-mediated endocytosis was inhibited (Figure 6E-F).

Taken together, our results from two targets (BRD4 and AR) and three E3 ligases (VHL, MDM2, and CRBN) suggested that CME is broadly required for PROTACs driven target degradation. This mechanism is summarized in our proposed model, where binding of PROTACs to CD36 leads to recruitment of PROTACs to clathrin coated pits. Internalization of PROTACs through the clathrin pathway is required for their ultimate release into the cellular interior, where they drive degradation of their targets (Figure 6G). Although the mechanisms governing PROTAC transit from clathrin coated pits to their cytosolic targets remains to be elucidated, our results point to a broad dependence on clathrin-mediated endocytosis for their entry into cells.

## Discussion

The involvement of CD36 in metabolic disease and cancer has long been recognized^9,68^. More recently, CD36 has been shown to facilitate the cellular uptake and activity of several PROTACs, extending its relevance to therapeutic delivery^14^. Given the physiological and therapeutic importance of its ligands, understanding how cells internalize CD36 is important for determining how these molecules are subsequently trafficked, processed, and utilized.

However, the endocytic route responsible for internalizing CD36 has itself remained unclear. Specifically, the long-standing assumption that CD36 is internalized by caveolae^19–21^ has recently been challenged by proteomic screening data that suggested components of clathrin-mediated endocytosis associate with CD36^14^. Therefore, we set out to evaluate the extent to which cellular uptake of CD36 and associated PROTACs requires the clathrin pathway. Our quantitative live cell imaging results illustrate that CD36 localizes strongly to sites of clathrin-mediated endocytosis but does not exceed nonspecific background levels of colocalization with caveolae. Localization to sites of clathrin-mediate endocytosis was progressively lost through K to R and Y to A mutations of residues within the CD36 cytosolic C-terminal domain, suggesting that recruitment of CD36 relies on multi-valent interaction with adaptor proteins of the clathrin pathway that recognize ubiquitin, such as Eps15^69,70^ and Epsin^71–73^, and those that recognize tyrosine-containing motifs, such as AP2^74^. Further we found that both the natural and therapeutic ligands of CD36, stearic acid and PROTACs, respectively, concentrate at endocytic sites in a CD36-dependent manner, and that accumulation of ligands within cells can be prevented by the inhibition of CME. Finally, we demonstrate that PROTACs with distinct chemistries, E3 ligase interactions, and target proteins all fail to degrade their targets when clathrin-mediated endocytosis is inhibited.

For fatty acid metabolism, uptake of long-chain fatty acids through CME suggest that recruitment to endocytic structures may contribute to the regulation of cellular fatty acid availability. Previous work has established a role for CD36 in tissue fatty acid uptake^75–77^. Our findings motivate further investigation into how the CD36 trafficking pathway influences the allocation of fatty acids to oxidation, storage, or membrane lipid synthesis. Because the present experiments used a fluorescent stearic acid analogue, extending these observations to unmodified fatty acids and additional metabolic cell types will be important.

Given that the requirement for clathrin-mediated endocytosis extends to multiple therapeutic degradation targets and E3 ligases, the kinetics of CME likely play an important role in determining PROTAC potency. This relationship is particularly relevant in cancer, where endocytic rates are frequently altered to support tumor progression^78,79^. For instance, cancer cells can selectively accelerate CME through a signaling-induced dynamin-1 pathway to modulate death receptor complex levels and evade apoptosis^80^. Since our data shows that PROTAC accumulation is effectively blocked when clathrin assembly or recruitment is inhibited, it follows that accelerated endocytosis in malignant cells could enhance the intracellular flux of PROTACs. This reasoning suggests that the high uptake efficiency of CD36-mediated degraders might be intrinsically linked to the upregulated endocytic kinetics characteristic of specific tumors.

Beyond the immediate implications for CD36, the finding that PROTAC uptake requires clathrin-mediated endocytosis suggests that bifunctional degraders could be engineered to hijack receptors other than CD36 that undergo robust clathrin-mediated internalization, such as the transferrin receptor^42,81^, receptor tyrosine kinases^82,83^, G protein-coupled receptors^84,85^, or immunological receptors^86,87^. By engineering the targeting ligand to bind receptors that are highly expressed or rapidly internalized in specific cell types, PROTACs could achieve superior tissue selectivity and intracellular delivery.

## Materials and Methods

### Cell culture and transfection

The cells used in this study were human SUM159 cells genetically edited to express HaloTag on both alleles of the AP2-σ2 (a gift from T. Kirchhausen). Cells were cultured in 1:1 high-glucose DMEM: Ham’s F-12 medium (Hyclone, GE Healthcare) supplemented with 5% fetal bovine serum (Hyclone), penicillin/ streptomycin/l-glutamine (Hyclone), 1 μg/ml hydrocortisone (H4001; Sigma-Aldrich), 5 μg/ml insulin (I6634; Sigma-Aldrich), and 10 mM HEPES buffer (pH 7.4). Culture conditions were maintained at 37°C with 5% CO₂. Cover slips used for cell seeding were acid-washed and heat-treated to ensure surface cleanliness and cell adhesion. Cover slips were placed in 6-well plates, with 30000 cells seeded per well. After overnight seeding, cells were transfected using Fugene HD transfection reagent (Promega) at a ratio of 3 μL Fugene HD per 1 μg plasmid DNA. After 24h post-transfection, cells were treated with 100 nM JF_646_ HaloTag ligand to label AP2-*σ*2 on the cell membrane surface and incubated at 37°C for another 15 min. Then cells were washed with warmed PBS and imaged immediately.

### Plasmid constructs

The mCherry-CD36 plasmid, a gift from Michael Davidson (Addgene #55011), served as the template for constructing three CD36 C-terminal mutants via site-directed mutagenesis. Specifically, the Ub-mut double mutant was generated by substituting lysine residues at positions 469 and 472 with arginines (K469R/K472R), while the Tyr-mut CD36 point mutant was engineered by replacing the tyrosine at position 463 with alanine (Y463A). Additionally, a triple-mutant construct, designated as Tyr/Ub-mut CD36, was created to incorporate all three aforementioned substitutions (Y463A/K469R/K472R) within the C-terminal tail.

The expression plasmid for (YTRF) TfRΔecto-mRFP was engineered by substituting the GFP domain in our previously reported (YTRF) TfRΔecto-GFP construct with mRFP^44^. This original plasmid encodes the human transferrin receptor (TfR) intracellular and transmembrane domains (residues 1–88; GenBank AAA61153) fused to GFP via a 9-amino acid flexible linker (GKGDPPVAT). The mRFP sequence was PCR-amplified from the pcDNA3-mRFP vector, which was kindly provided by Dr. Douglas Golenbock (Addgene #13032).

To generate the (CTRD) TfR-Δecto-mRFP variant, which exhibits diminished affinity for sites of clathrin-mediated endocytosis), the YTRF motif at positions 20-23 within the TfR intracellular domain was converted to CTRD using sequential site-directed mutagenesis. In the first step, position 20 was mutated to cysteine to yield a (CTRF) TfR-Δecto-mRFP intermediate. Subsequently, Position 23 was substituted with aspartic acid to produce the final (CTRD) model receptor variant.

The fusion construct, AP180ct-mCherry, was synthesized by GenScript using the FLASH Gene Synthesis service. The cDNA sequence encoding the C-terminal fragment of AP180 (AP180ct, residues 516 to 898) was fused in frame with a C-terminal mCherry tag. The synthetic fragment was codon-optimized for mammalian expression and subcloned into the pcDNA3.1(+)-N-Myc expression vector between the *EcoRI* and *NotI* restriction sites. This strategy resulted in a fusion protein containing an N-terminal Myc tag, followed by AP180ct and a C-terminal mCherry. Caveolin-1–GFP (Addgene plasmid #14433), caveolin1-mCherry (Addgene plasmid #27705), and cavin1-EGFP (Addgene plasmid #27709) were purchased from Addgene. The cavin1-RFP construct was generated by GenScript by replacing the EGFP coding sequence in the cavin1-mEGFP plasmid with that encoding RFP.

All plasmid constructs were custom-synthesized by GenScript. Following delivery, the integrity of each construct was validated via whole-plasmid sequencing (Plasmidsaurus) to ensure correct insertion, proper reading frames, and the presence of targeted mutations.

### Fluorescence microscopy

Olympus spinning disk confocal microscopy system was used for dynamic observation of live cell plasma membranes. This system integrates a Yokogawa CSU-W1 SoRa scanning unit and is built upon the Olympus IX83 microscope platform, equipped with a 100× Plan-Apochromat oil immersion objective (NA 1.5). Fluorescence signals were captured using a Hamamatsu ORCA C13440-20CU CMOS camera. To maintain cell viability, samples were equipped with a heating device to stabilize coverslip temperature at 37 °C. All plasma membrane imaging was completed 24 hours post-transfection. To significantly reduce photobleaching during live-cell imaging, imaging medium was phenol red-free and supplemented with OxyFluor deoxy reagent at a 1:33 volume ratio. The laser excitation scheme was configured as follows: 488 nm excitation for Atto 488, 561 nm excitation for RFP/mCherry, and 640 nm excitation for the JF646 HaloTag ligand of AP2, respectively.

### siRNA transfection and CD36 knock down

Seed cells at a density of 30000 cells per well in a six-well plate. Incubate at 37 ℃ in a 5% CO_2_ incubator for 24 h until cells adhere. Perform siRNA transfection using Lipofectamine RNAiMAX transfection reagent (Thermo Fisher Scientific, Cat. No. 13778075) according to the manufacturer’s instructions. The transfection complex preparation procedure is as follows: Take 6.5 μL of Lipofectamine RNAiMAX and 5.5 μL of 10 μM siRNA stock solution (Santa Cruz, sc-29995). Dilute each into 100 μL of Opti-MEM reduced serum medium and pre-incubate separately at RT for 5 min. Slowly add the diluted siRNA solution to the Lipofectamine RNAiMAX diluent, mix thoroughly, and continue incubating at room temperature for 10 min to form the transfection complex. Before adding the complex, replace the stale medium in the six-well plate with fresh complete medium. Then, evenly add the transfection complex to the wells by pipetting dropwise and gently mix. To ensure knockdown efficiency, transfection was performed in two rounds, spaced 24 hours apart. Control cells were simultaneously transfected with non-targeting control siRNA (purchased from QIAGEN, Germantown, MD). Finally, cells were harvested on day 4 post-seeding for subsequent measurements.

### Internalization assay

Based on previously established protocols, the internalization of PROTAC and transferrin (Tf) via clathrin-mediated endocytosis was quantified using flow cytometry. Initially, both PROTAC and transferrin were conjugated with Atto-488. SUM cells were seeded in 6-well plates at a density of 30000 cells/well in a total volume of 2 mL. After 24 h of incubation, the culture media was aspirated, and cells were rinsed three times with PBS. Following a 30 min starvation period in 1 mL of pre-warmed serum-free media, cells were incubated with of 0.5 μM PROTACs or Tf (50 μg/mL) at designated time points. To remove extracellularly bound cargo, cells were treated with 500 μL of trypsin for two times at 37 °C, this wash step was performed twice to ensure the elimination of excess PROTAC or Tf. Subsequently, cells were fully detached by a final 5 min incubation with trypsin. The enzymatic reaction was quenched with 1 mL of complete media, and the resulting cell suspension was collected and resuspended in 300 μL of ice-cold PBS for immediate flow cytometric analysis.

Flow cytometry analysis was performed using the Guava easyCyte system (Millipore Sigma-Aldrich) featuring multi-wavelength lasers (405/488/532 nm) for excitation and detection. During sample preparation, cells were resuspended in PBS to yield a single-cell suspension, thereby minimizing the effects of clumping and non-specific binding. The system was calibrated according to a standardized protocol to ensure consistent detection accuracy and reproducibility. The median fluorescence intensity (MFI) of the fluorescent protein linked to each plasmid was measured using flow cytometry to assess expression levels^88^. Throughout acquisition, the flow rate was kept at 35 μL/min, with thresholds optimized to filter background interference. FSC and SSC parameters were employed to characterize cell size and complexity, while cell debris was excluded by gating on the scatter plot. Finally, FlowJo software was used for analysis of the raw data, with each experimental conclusion derived from three biological replicates (n = 3).

### Immunoblotting assays

Seed cells at a density of 30000 cells per well in a six-well plate. After 24 hours of culture, add PROTAC and incubate for 4 hours. Subsequently, cells were washed with PBS and collected. Whole-cell lysates were prepared using RIPA buffer (50 mM Tris-HCl pH 7.4, 250 mM NaCl, 0.1% SDS, 0.5% sodium deoxycholate, 1% Nonidet P-40, 1 mM DTT) to prepare whole-cell lysates. Target protein expression levels were detected using protein-specific antibodies. Immunoreactive bands were visualized and quantified using the Odyssey imaging system after incubation with fluorescently labeled secondary antibodies. Primary antibodies used in the experiments included: BRD4 (ab128874) and CD36 (ab252922) from Abcam (Cambridge, UK); AR (ab108341) from Santa Cruz Biotechnology; and the housekeeping control antibody GAPDH (Cat# 5174) from Cell Signaling Technology (Danvers, MA, USA).

### Image analysis

CMEAnalysis (Danuser Lab) was employed to identify clathrin-coated structures and quantify their intensities appearing as fluorescent puncta in confocal images^47^. Throughout the imaging processing workflow, the AP2-σ2 signal served as the master channel for tracking these clathrin-coated structures. Two-dimensional Gaussian functions were utilized to fit local intensity maxima in the mCherry/RFP channels. The Gaussian standard deviation was derived from the physical parameters of the microscope to estimate the point spread function. To ensure the accuracy of the fit, residuals were subjected to Anderson-Darling testing. The fluorescence intensity (corresponding to Gaussian amplitude) and spatial position of each detected puncta were recorded. As established in prior studies, candidate puncta must exhibit diffraction-limited characteristics and possess brightness significantly higher than surrounding membrane components to be classified as valid clathrin-coated structures^48,89^. For validated puncta in the master channel, two-dimensional Gaussian fitting was simultaneously performed on corresponding transmembrane fusion protein, receptor, or ligand channels within a region centered at the spot’s position and having a radius of 3σ pixels^48^.

### Statistical analysis

Micrograph acquisition experiments were independently repeated on separate days, producing consistent results across replicates. For CMEAnalysis, cell image data were collected independently under at least two different experimental conditions or on separate days. As described throughout this paper, all reported experimental results are based on a minimum of three independent replicates. For comparisons with statistically significant differences, a two-sample Student’s *t*-test assuming unequal variances was used. The resulting p-values are provided in the corresponding figure captions where such comparisons are presented.

## Data, Materials and Software Availability

All study data are included in the article and/or *SI Appendix*.

## Acknowledgements

We thank T. Kirchhausen for the gift of SUM159/AP2-σ2 Halo Tag cells. This research was supported by the National Institutes of Health through grants R35GM139531 to J.C.S., R01CA277682 to H.Y. Li, RO1AI096090 to J.M.H., and the Welch Foundation through grant F-2257 to J.C.S.

## Author contributions

H.-Y. Liu and J.C.S. designed the experiments. H.-Y. Liu and R. S. designed the plasmids. H.-Y. Liu performed live-cell imaging experiments. H.-Y. Li, Z. W., H.Z, and M. W. designed and synthesized PROTAC molecules. H.-Y. L. conducted western blot with the help of R. S., J. P. and B.K. H.-Y. Liu and J.C.S. wrote the manuscript, with J.M.H. and H.-Y. Li provided consultation on manuscript preparation and editing. H.-Y. Li also provided early guidance for the project. All authors reviewed and approved the final version of the manuscript.

## Supplementary Materials

**Figure S1.**
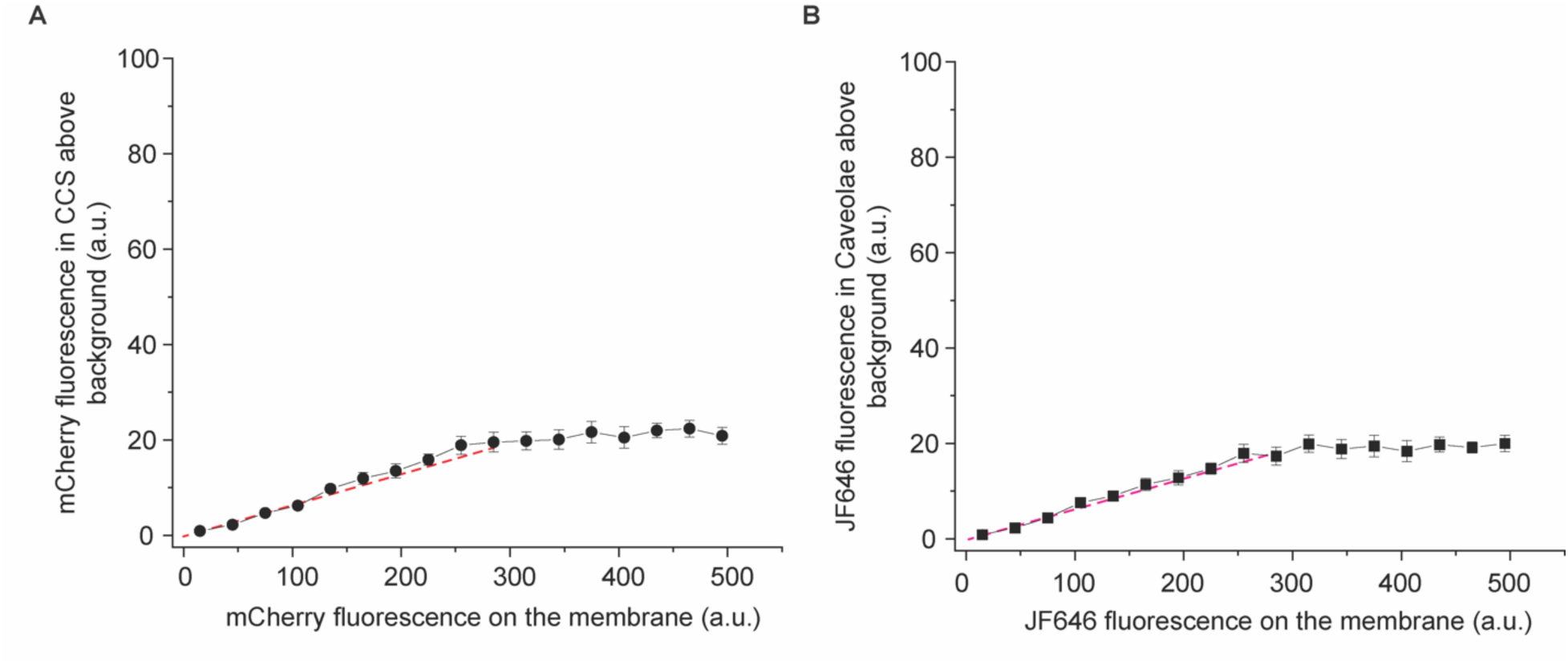
Quantification of caveolin1-mCherry and AP2-JF646. (A) Using AP2-JF646 as the main channel to identify clathrin-coated structures (CCSs), caveolin1-mCherry fluorescence at CCSs above local background is plotted against its fluorescence intensity in the surrounding plasma membrane. (B) Using caveolin1-mCherry as the main channel to identify caveolae, AP2-JF646 fluorescence at caveolae above local background is plotted against its fluorescence intensity in the surrounding plasma membrane. The red dashed line represents a linear fit to the initial linear region of the curve, and indicates the level of correlation expected from incidental colocalization. The similar slopes are consistent with incidental colocalization in both analyses. To provide a conservative estimate of incidental colocalization, the fit in panel A, which has the larger slope, was used as the reference in the main text analyses. Data were collected from over 5000 endocytic sites respectively. Data represent mean ± s.e.m..

**Figure S2.**
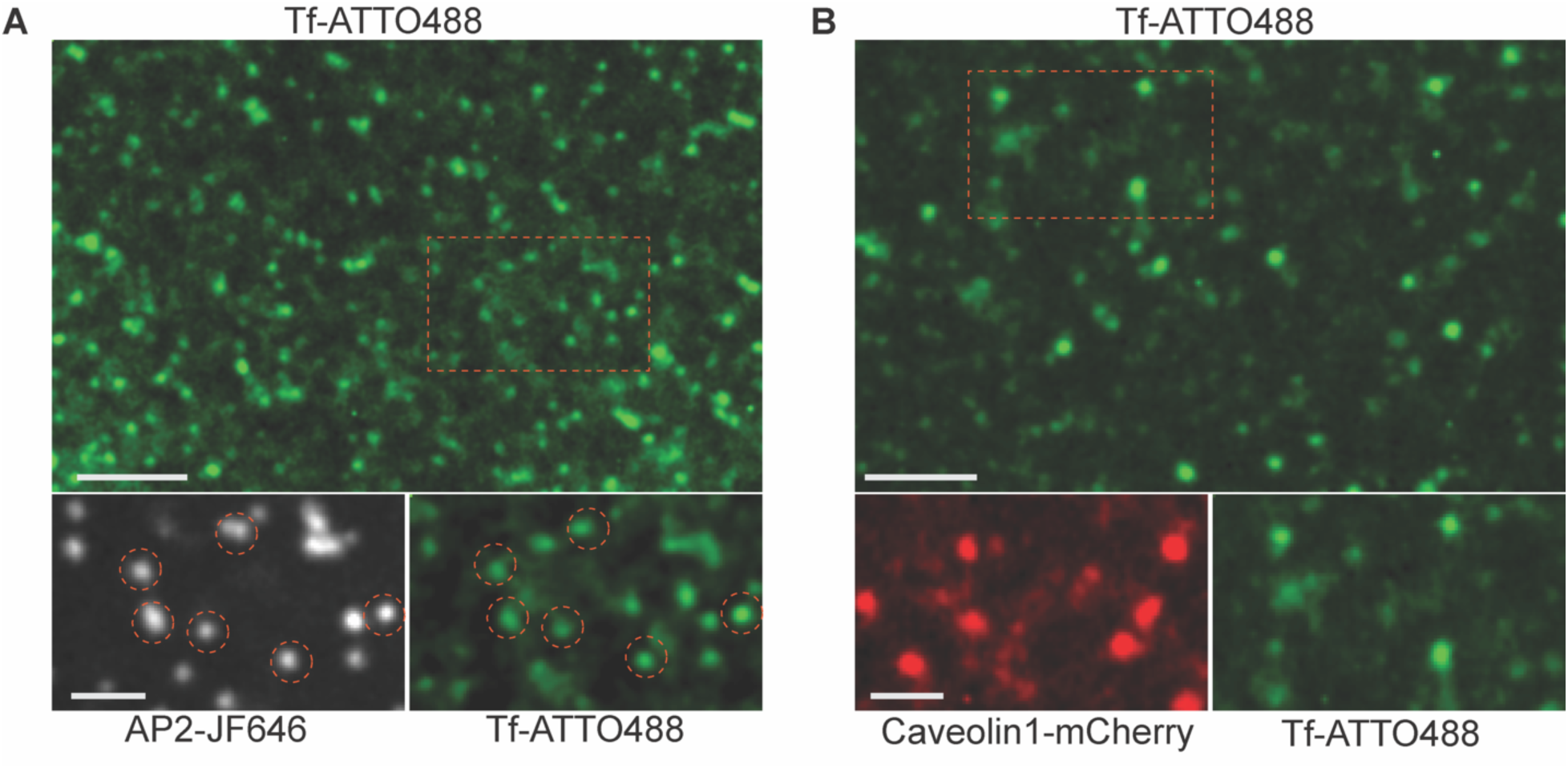
Transferrin (Tf) selectively colocalizes with AP2-positive clathrin-coated structures but not caveolae. (A) Representative fluorescence images showing the colocalization of Tf-ATTO488 (20 ng/µL, green) with the clathrin adaptor AP2-JF646 (grey). Red dashed circles in the inset demonstrate distinct recruitment and overlap of Tf-ATTO488 at AP2-positive endocytic sites. (B) Representative fluorescence images of cells expressing the caveolar marker Caveolin1-mCherry (red) and treated with Tf-ATTO488 (20 ng/µL, green). Tf-ATTO488 shows almost no significant colocalization with Cav1-positive caveolae. Scale bars, 5 µm (main panels) and 2 µm (insets).

**Figure S3.**
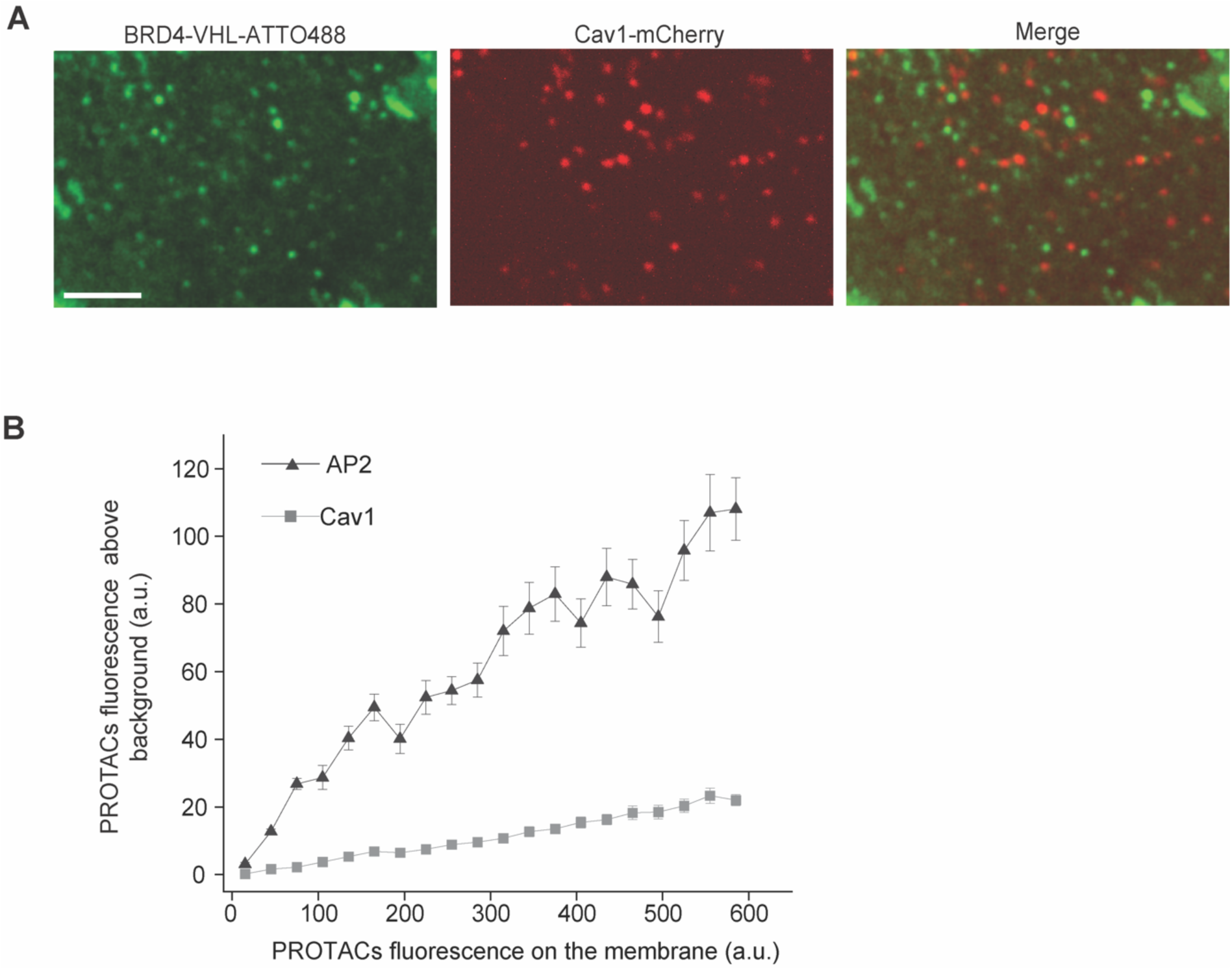
PROTACs are selectively excluded from caveolae. (A) Representative spinning disk confocal images of SUM 159 cells expressing the caveolar marker Cav1-mCherry (red) and treated with the fluorescent PROTAC, BRD4-VHL-ATTO488 (green). The merged image demonstrates a distinct lack of colocalization, indicating that PROTAC molecules do not partition into caveolar domains. Scale bar, 5 µm. (B) Quantification of PROTAC recruitment to clathrin-coated structures (AP2) versus caveolae (Cav1). The fluorescence intensity of PROTACs within each respective endocytic structure is plotted against the local PROTAC fluorescence on the plasma membrane. While PROTACs exhibit robust, concentration-dependent recruitment to AP2-positive sites, their accumulation within Cav1-positive caveolae remains negligible. Data represent mean ± SEM.

**Figure S4.**
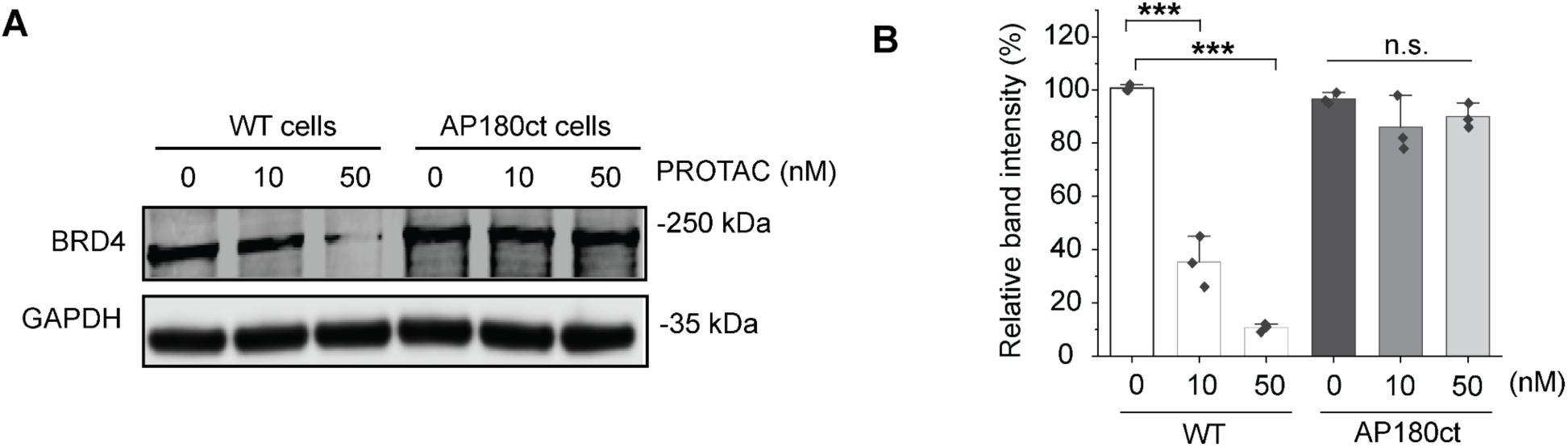
Genetic inhibition of CME abolishes PROTAC-mediated protein degradation. (A, B) Representative Western blots (A) and corresponding quantitative analysis (B) showing BRD4 protein levels in wild-type (WT) cells and cells expressing AP180ct (a dominant-negative inhibitor of CME) treated with indicated concentrations of PROTAC (0, 10, 50 nM). Expression of AP180ct significantly blocked the degradation of BRD4 induced by the PROTAC. Band intensities were normalized to GAPDH. Data are presented as mean ± SD from three independent experiments. *** P < 0.001, n.s., no significant difference.

**Scheme S1.**
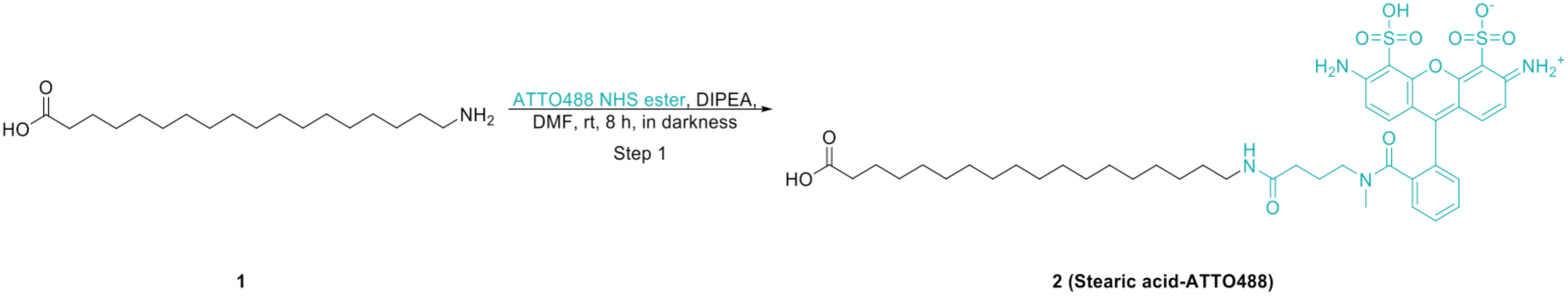
Reaction scheme for the synthesis of Stearic acid-ATTO488.

**Scheme S2.**
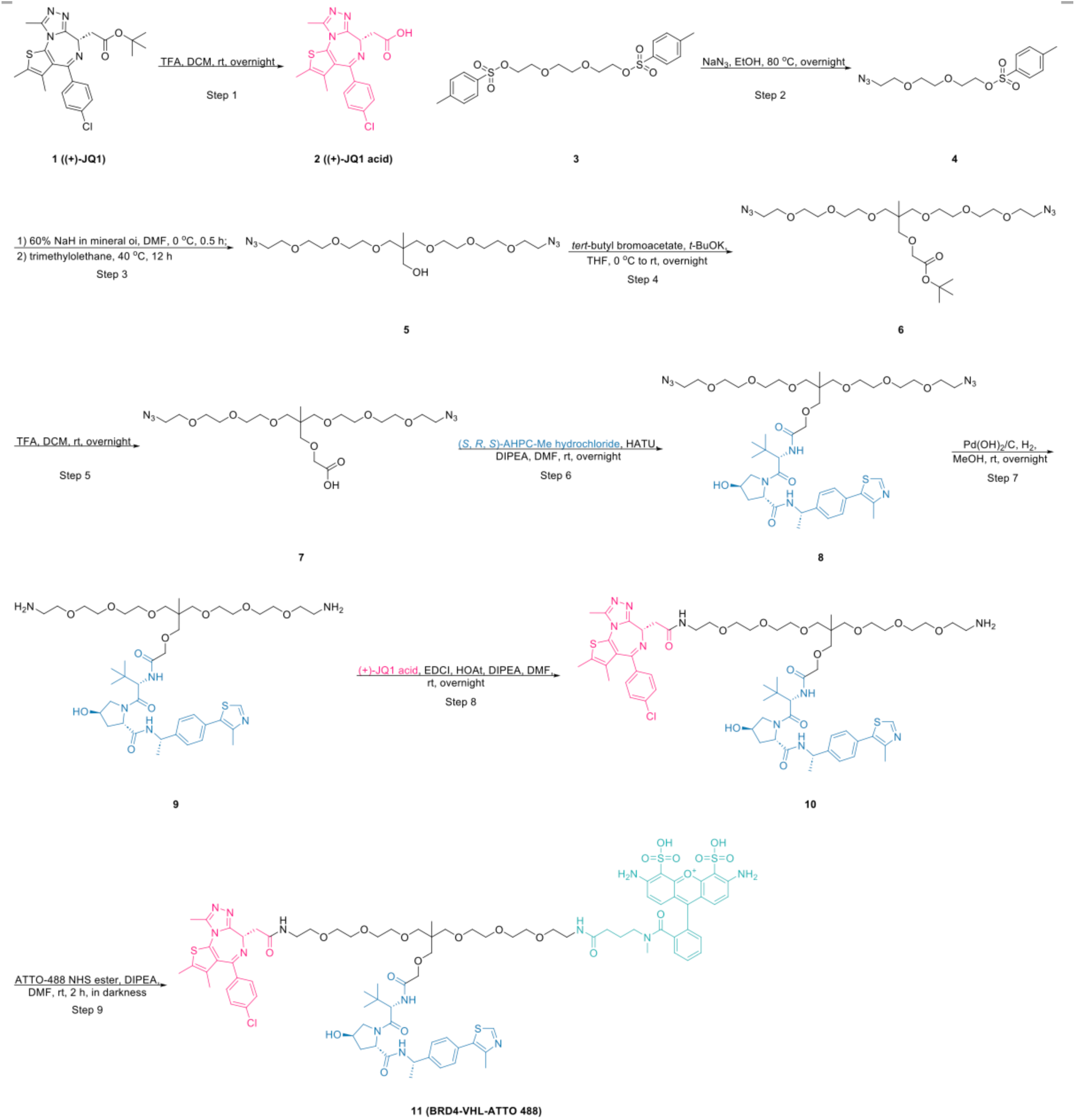
Reaction scheme for the synthesis of BRD4-VHL-ATTO488.

**Figure S5.**
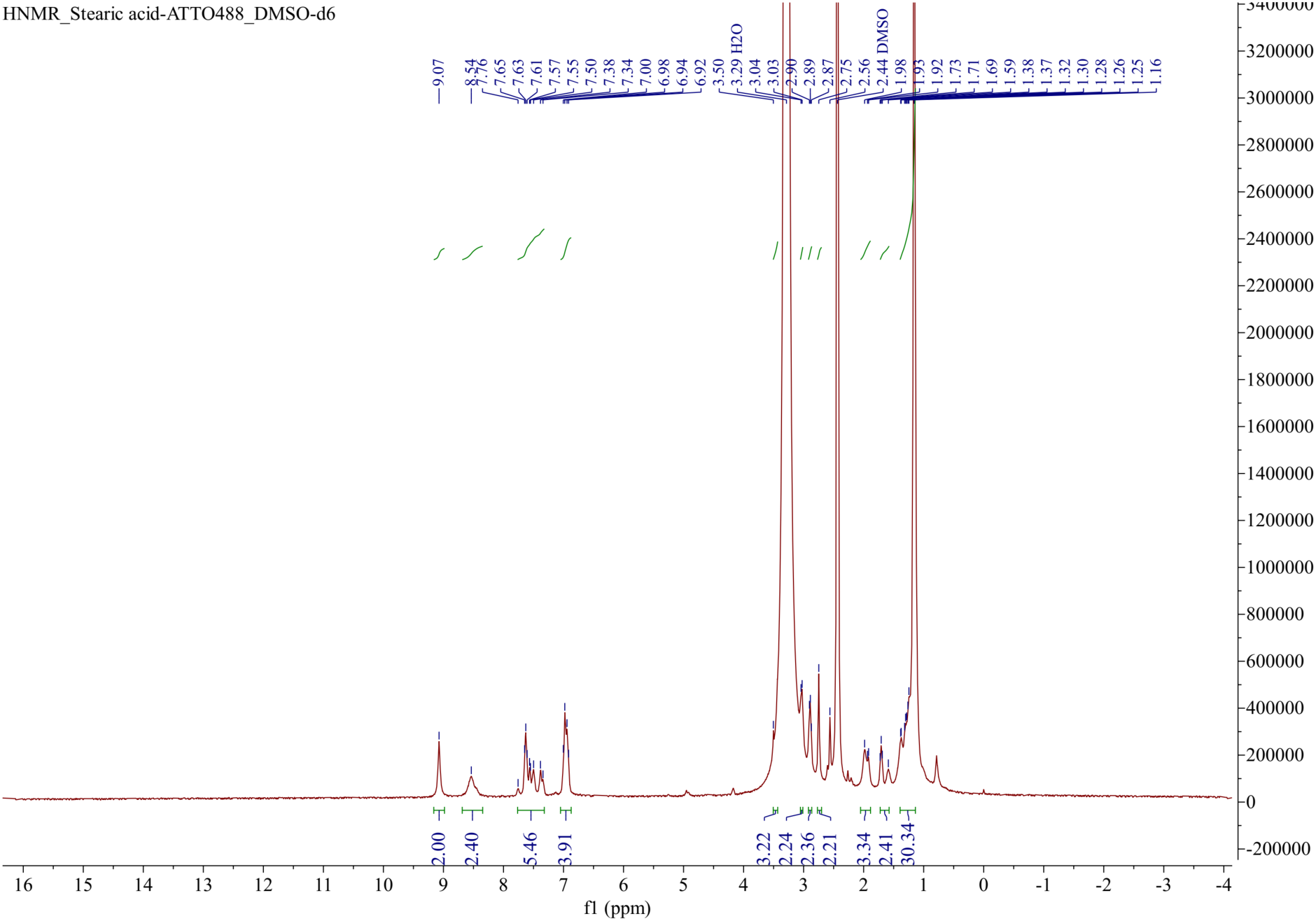
1H NMR spectrum (400 MHz, DMSO-d6) of Stearic acid-ATTO488.

**Figure S6.**
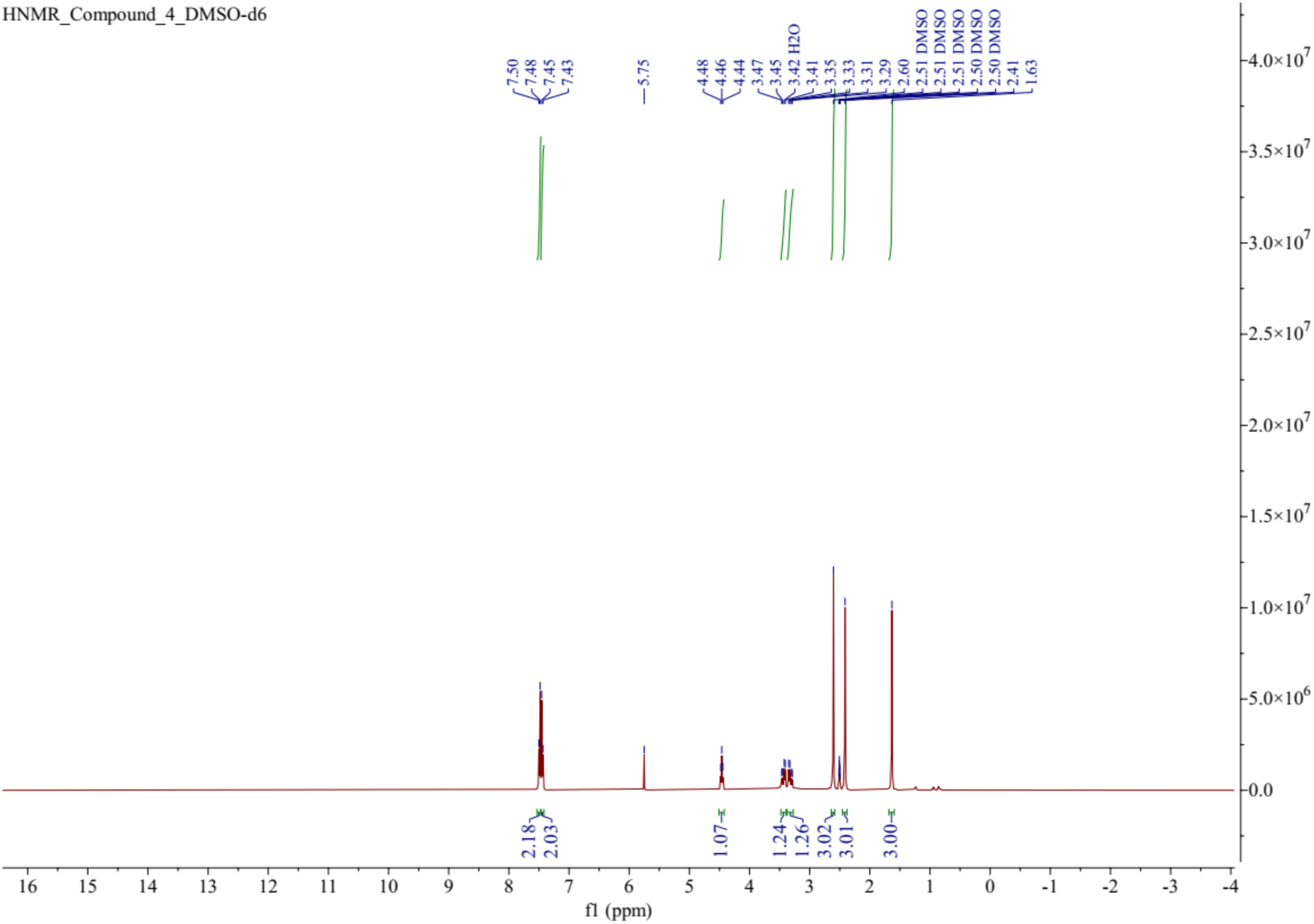
1H NMR spectrum (400 MHz, DMSO-d6) of Compound 4.

**Figure S7.**
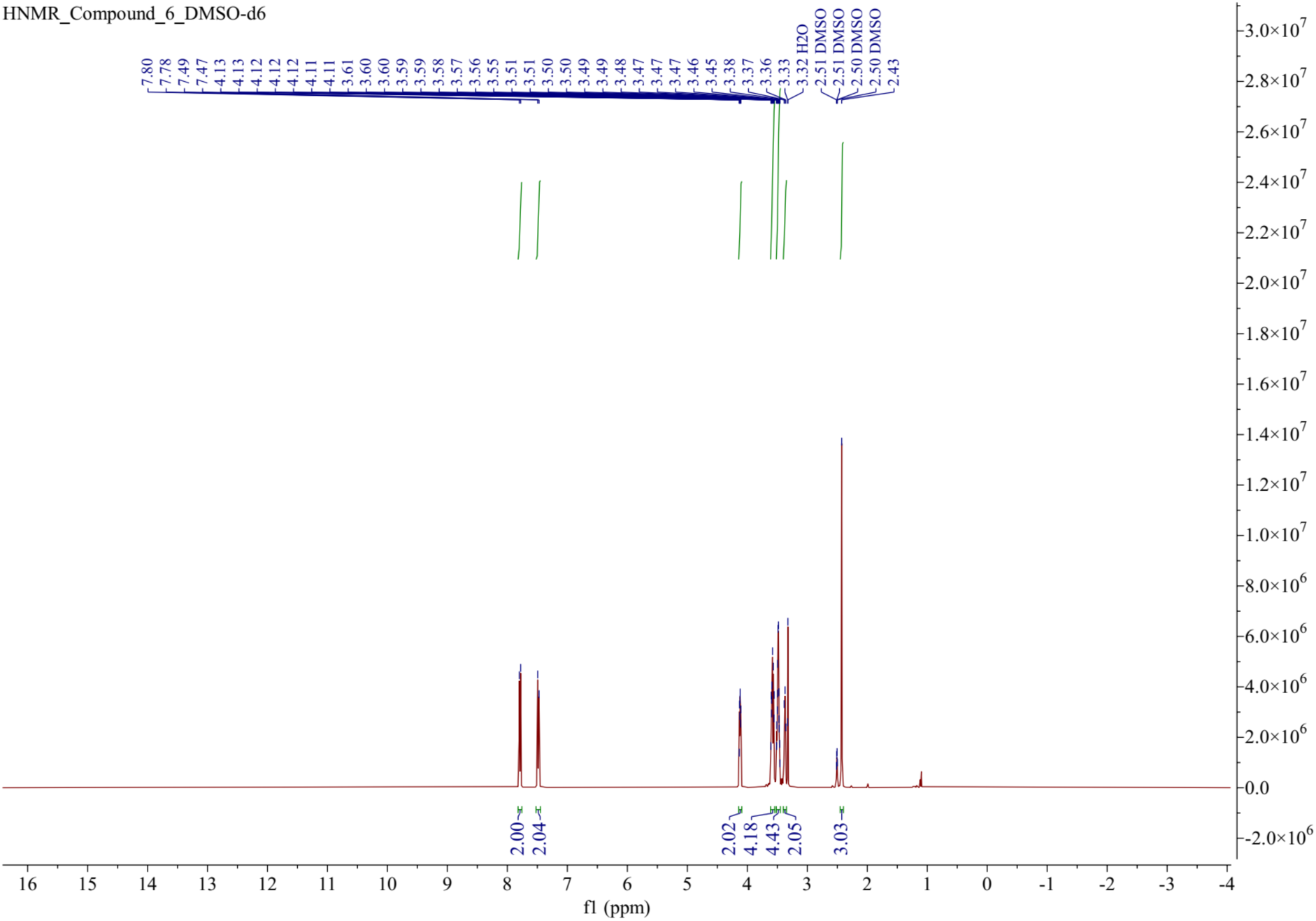
^1^H NMR spectrum (400 MHz, DMSO-*d6*) of Compound 6.

**Figure S8.**
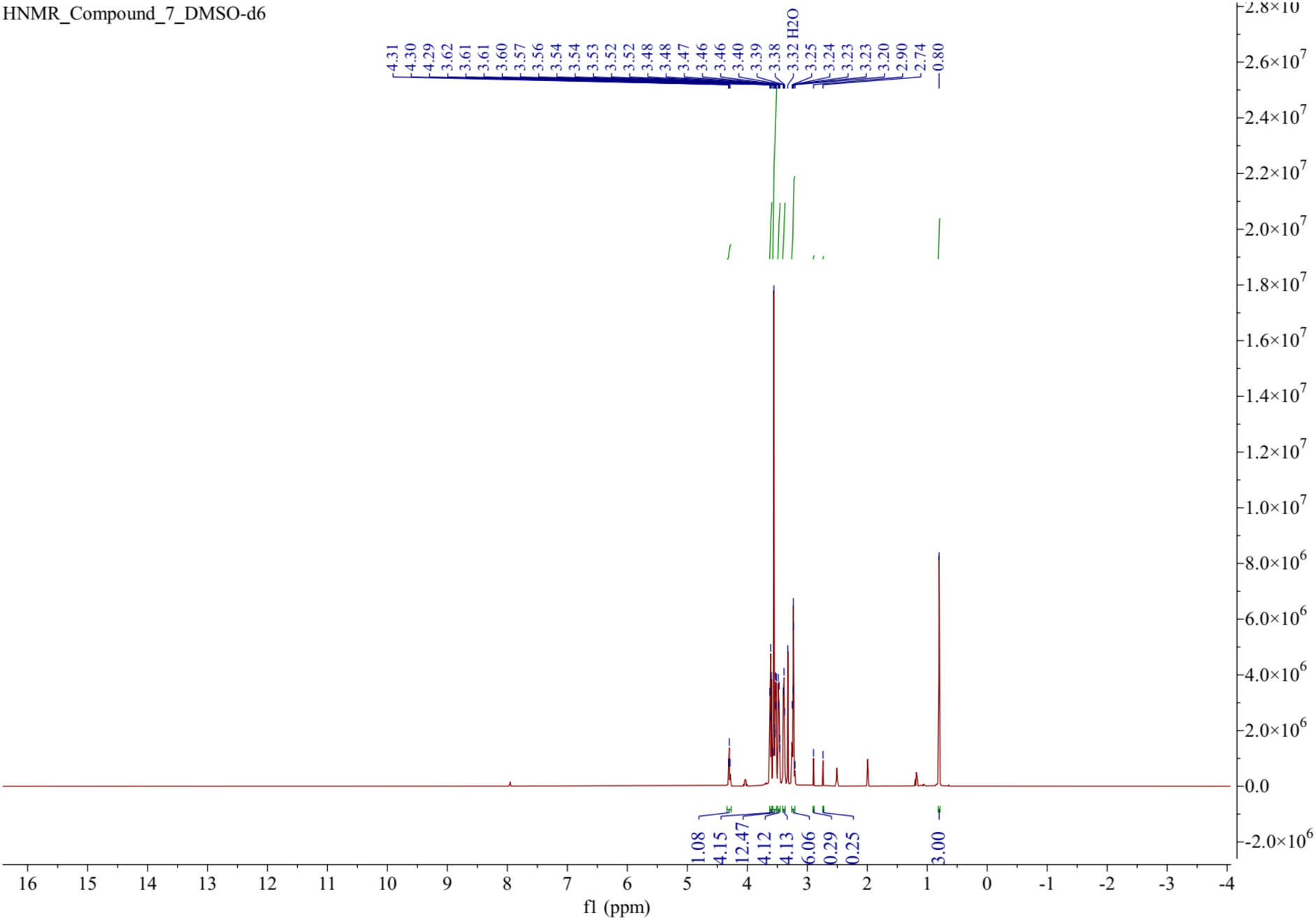
^1^H NMR spectrum (400 MHz, DMSO-*d6*) of Compound 7.

**Figure S9.**
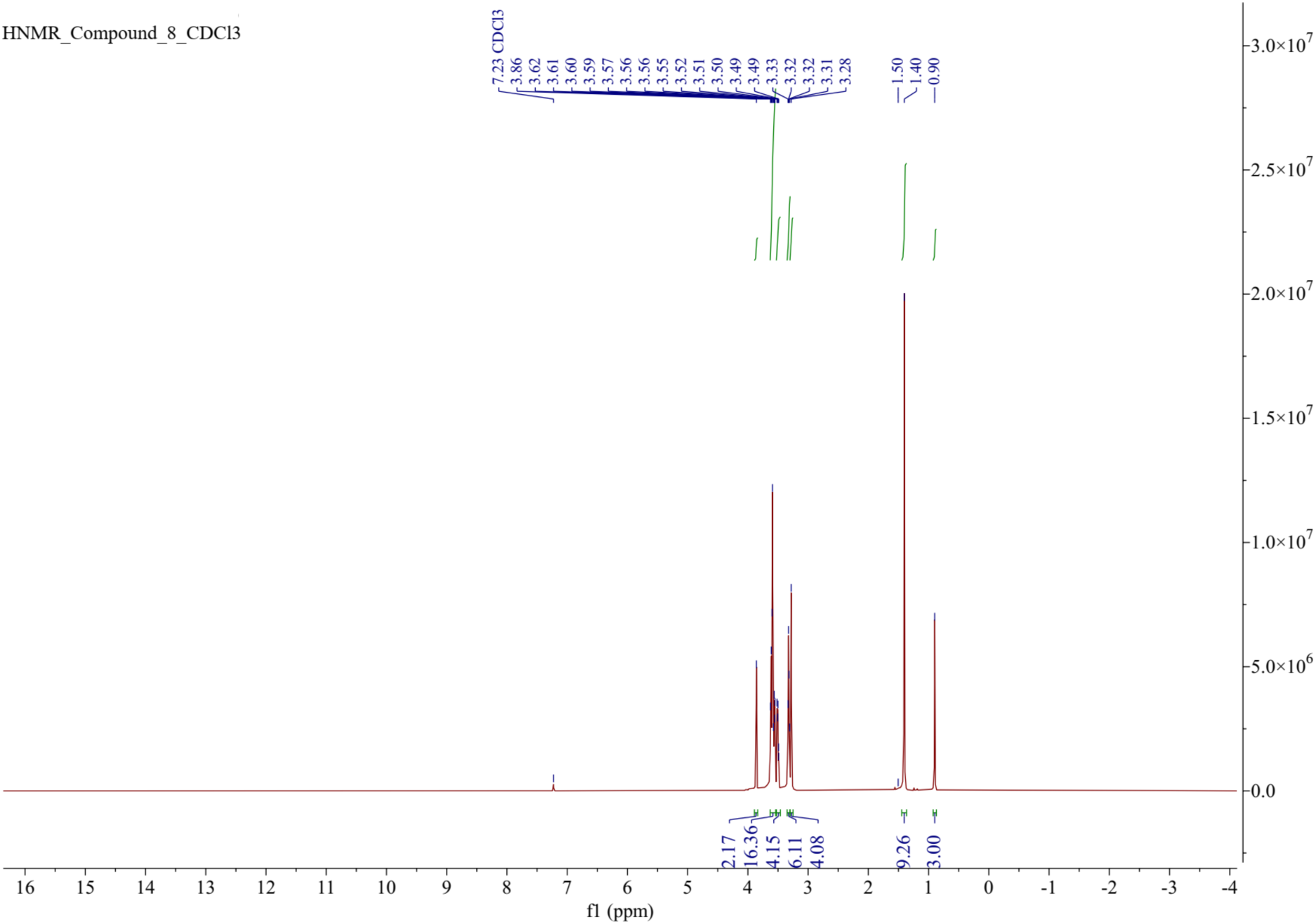
^1^H NMR spectrum (400 MHz, DMSO-*d6*) of Compound 8.

**Figure S10.**
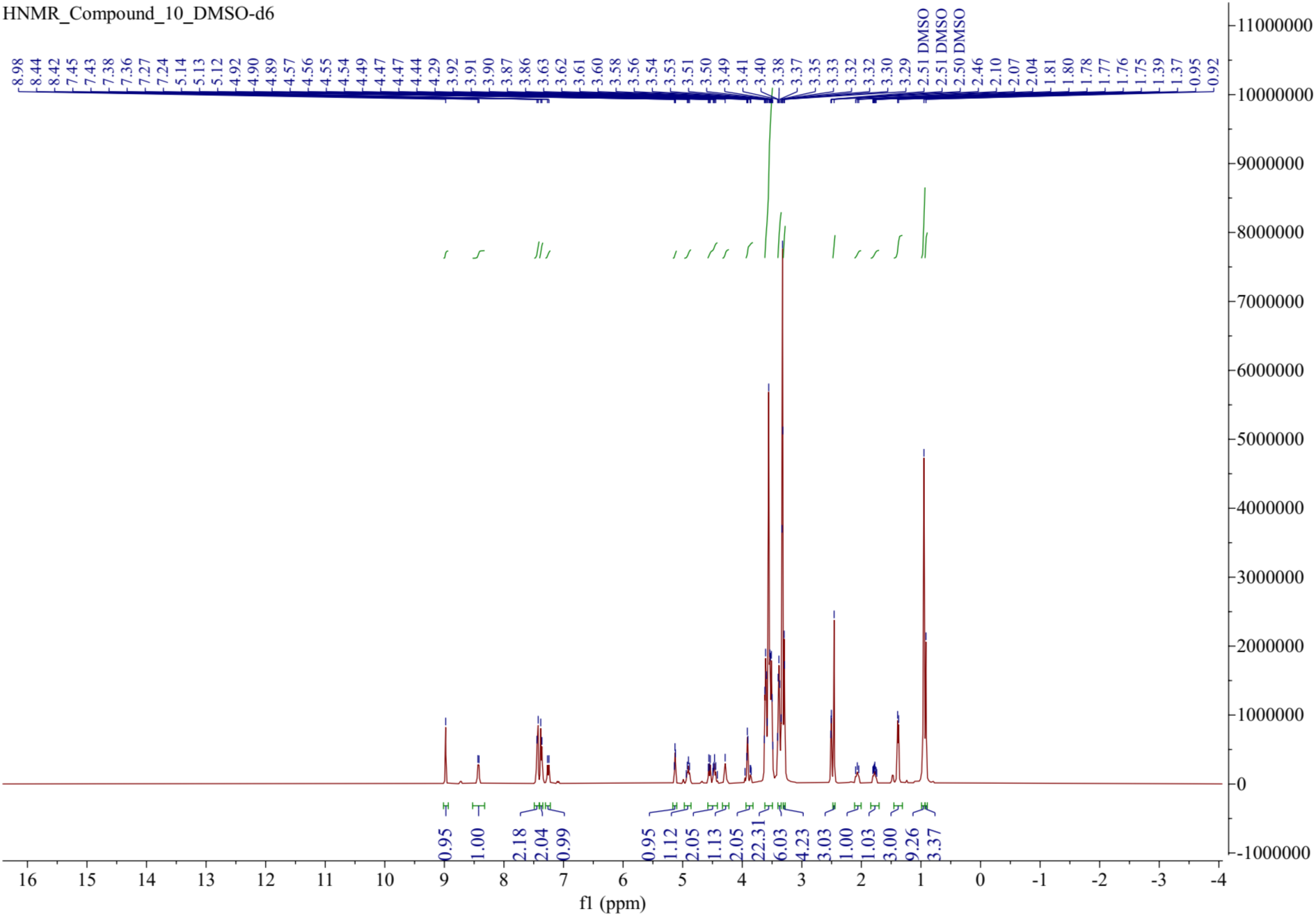
^1^H NMR spectrum (400 MHz, CDCl_3_) of Compound 10.

**Figure S11.**
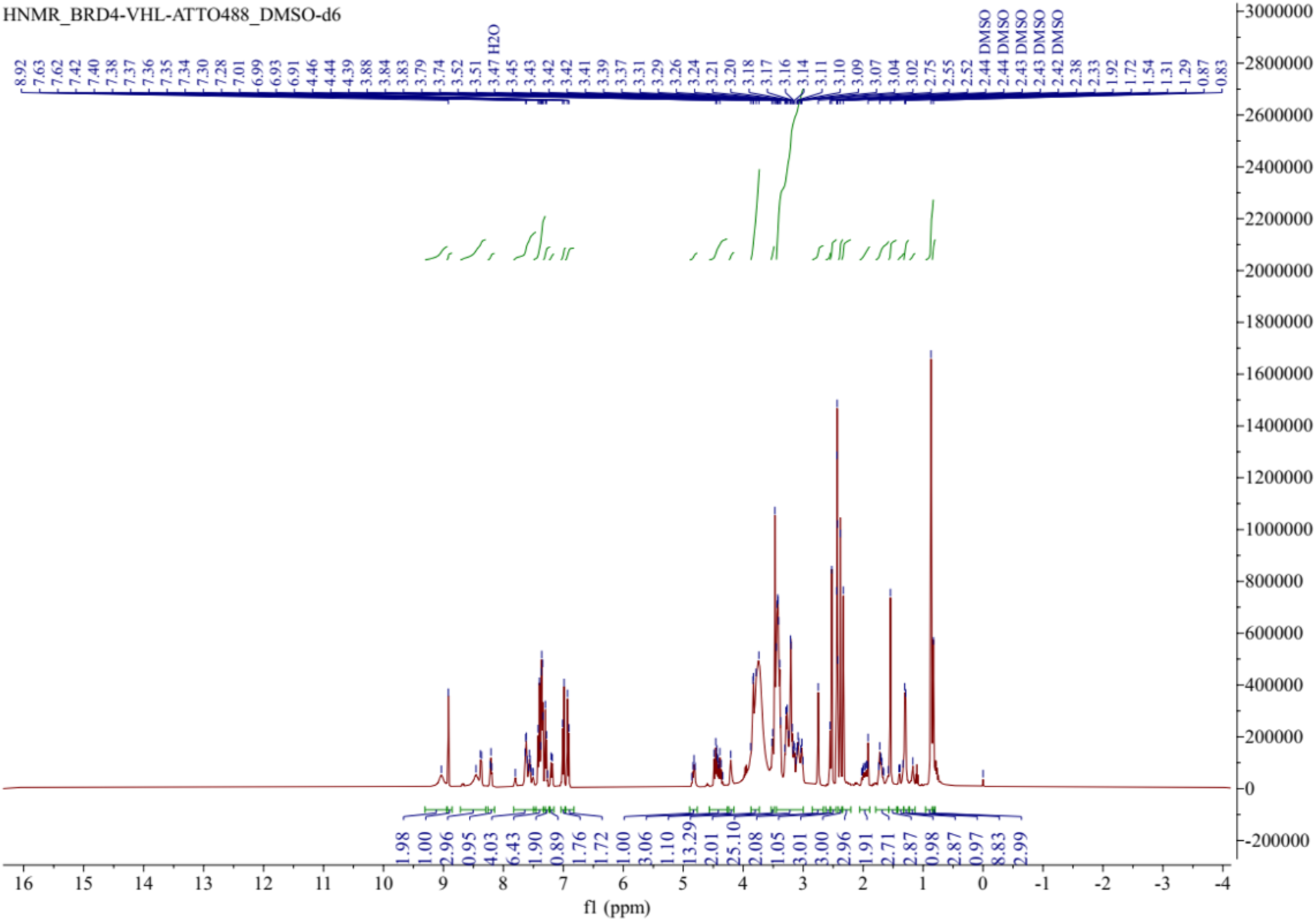
^1^H NMR spectrum (400 MHz, DMSO-*d6*) of BRD4-VHL-ATTO488.

**Figure S12.**
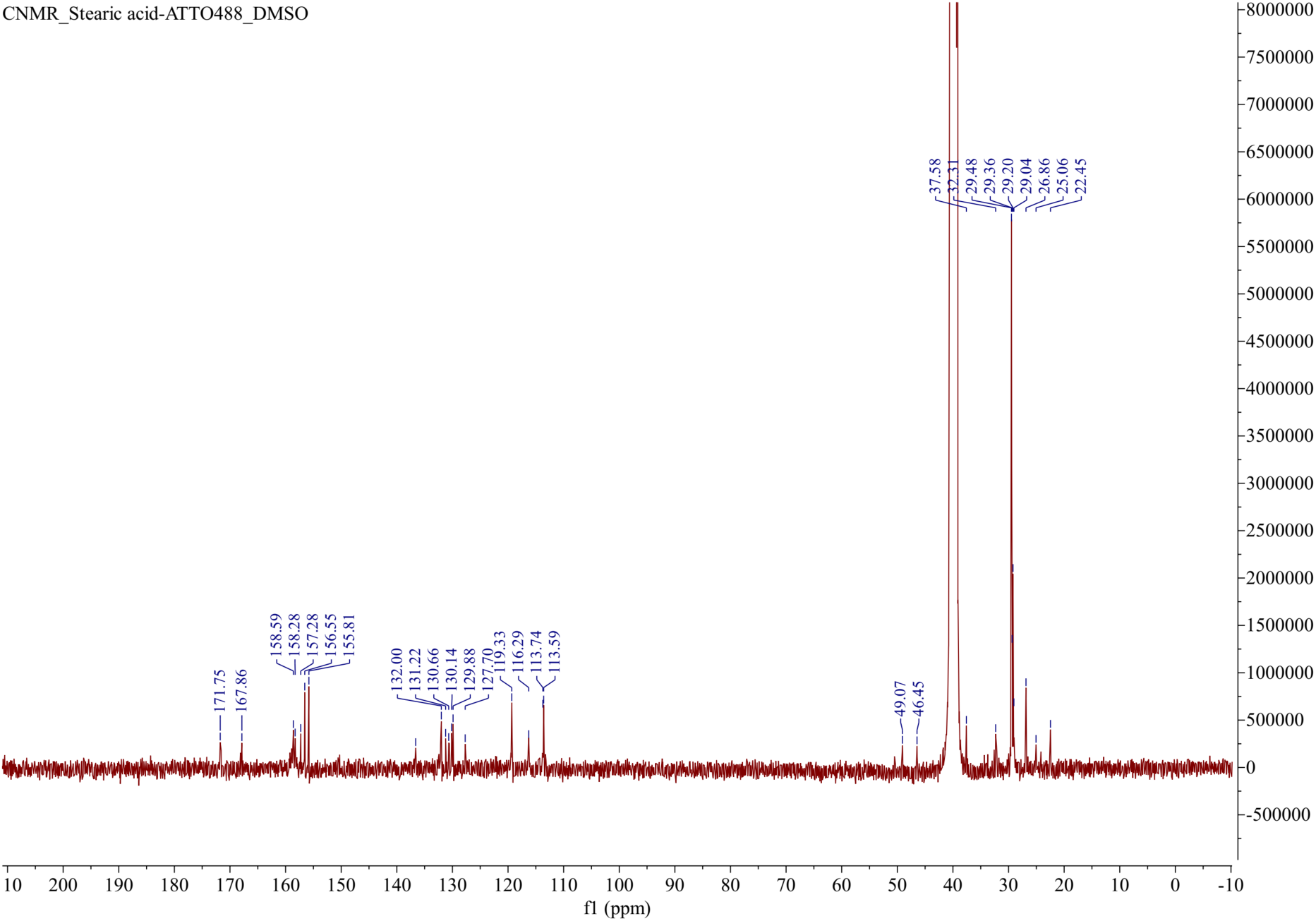
^13^C NMR spectrum (101 MHz, DMSO-d6) of Stearic acid-ATTO488.

**Figure S13.**
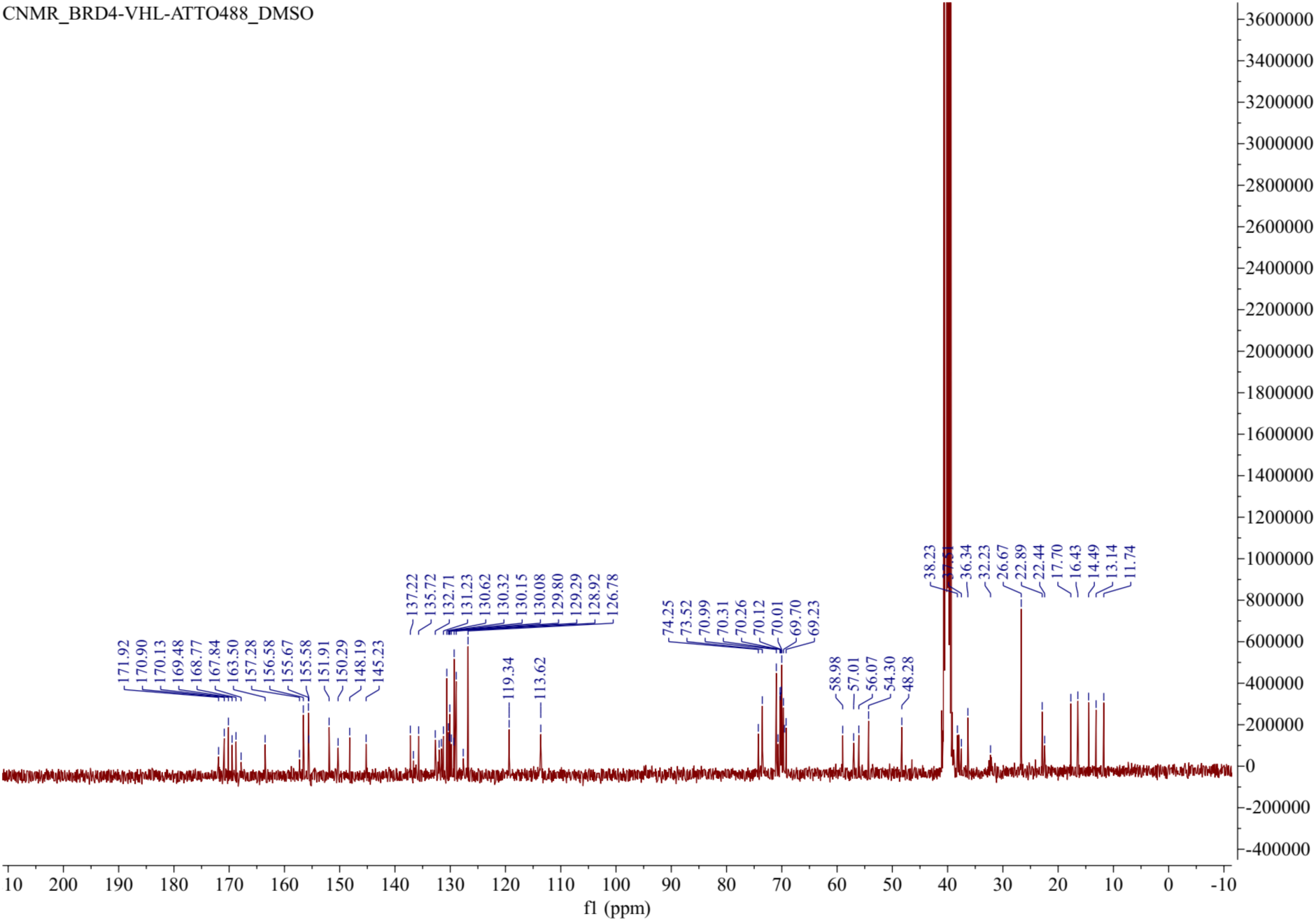
^13^C NMR spectrum (101 MHz, DMSO-d6) of BRD4-VHL-ATTO488.

**Figure S14.**
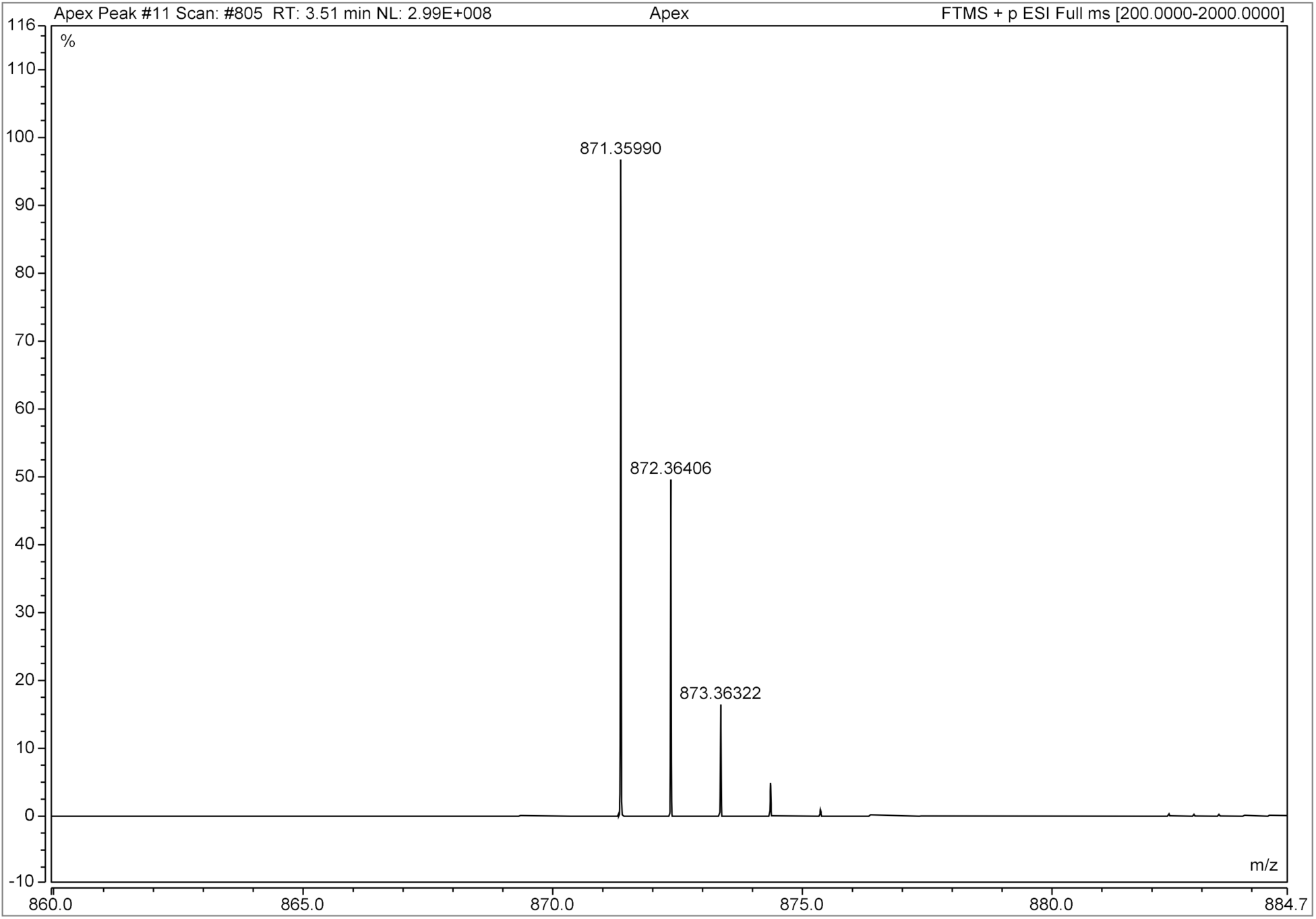
HR-MS spectrum of Stearic acid-ATTO488.

**Figure S15.**
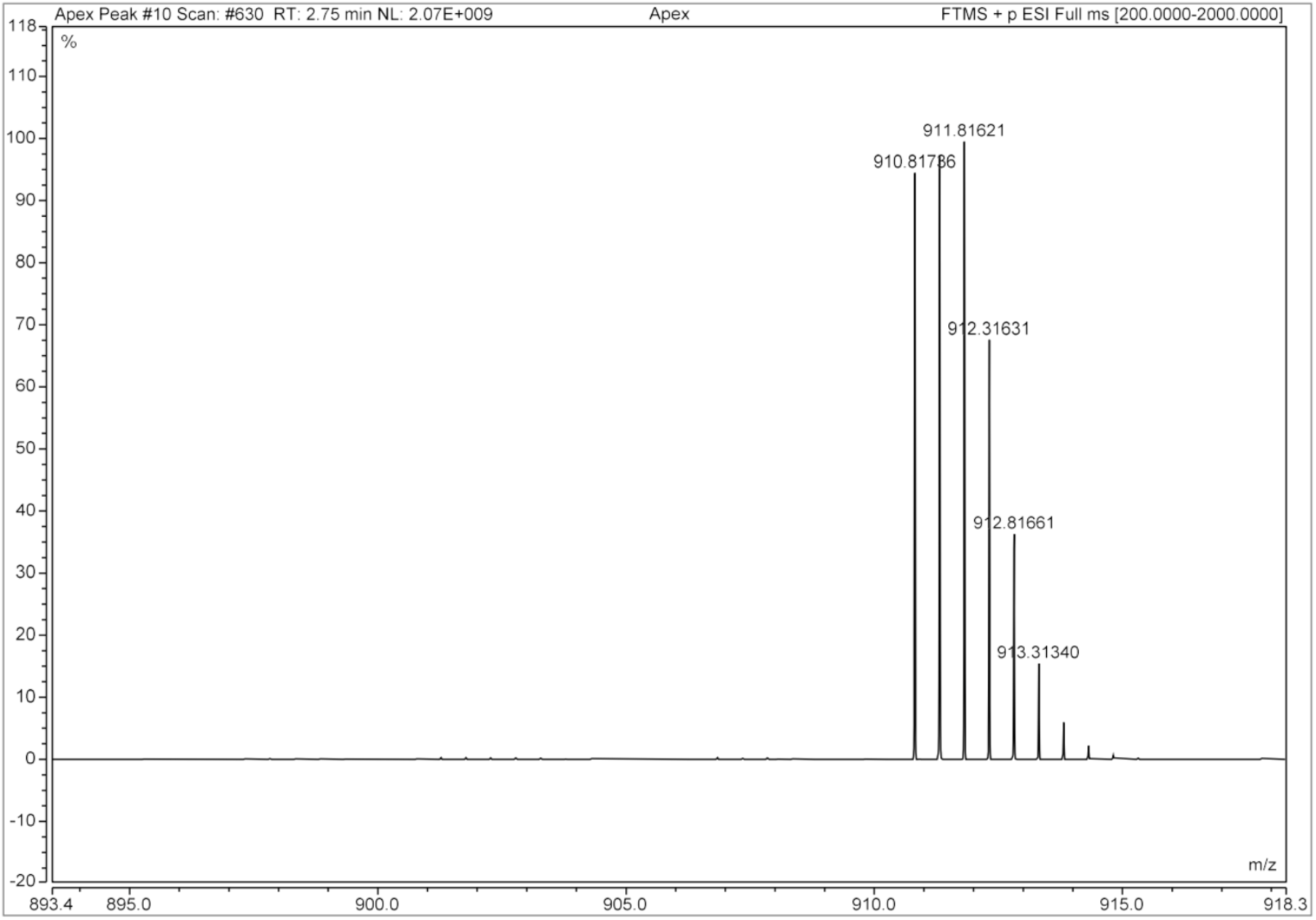
HR-MS spectrum of BRD4-VHL-ATTO488.

**Figure S16.**
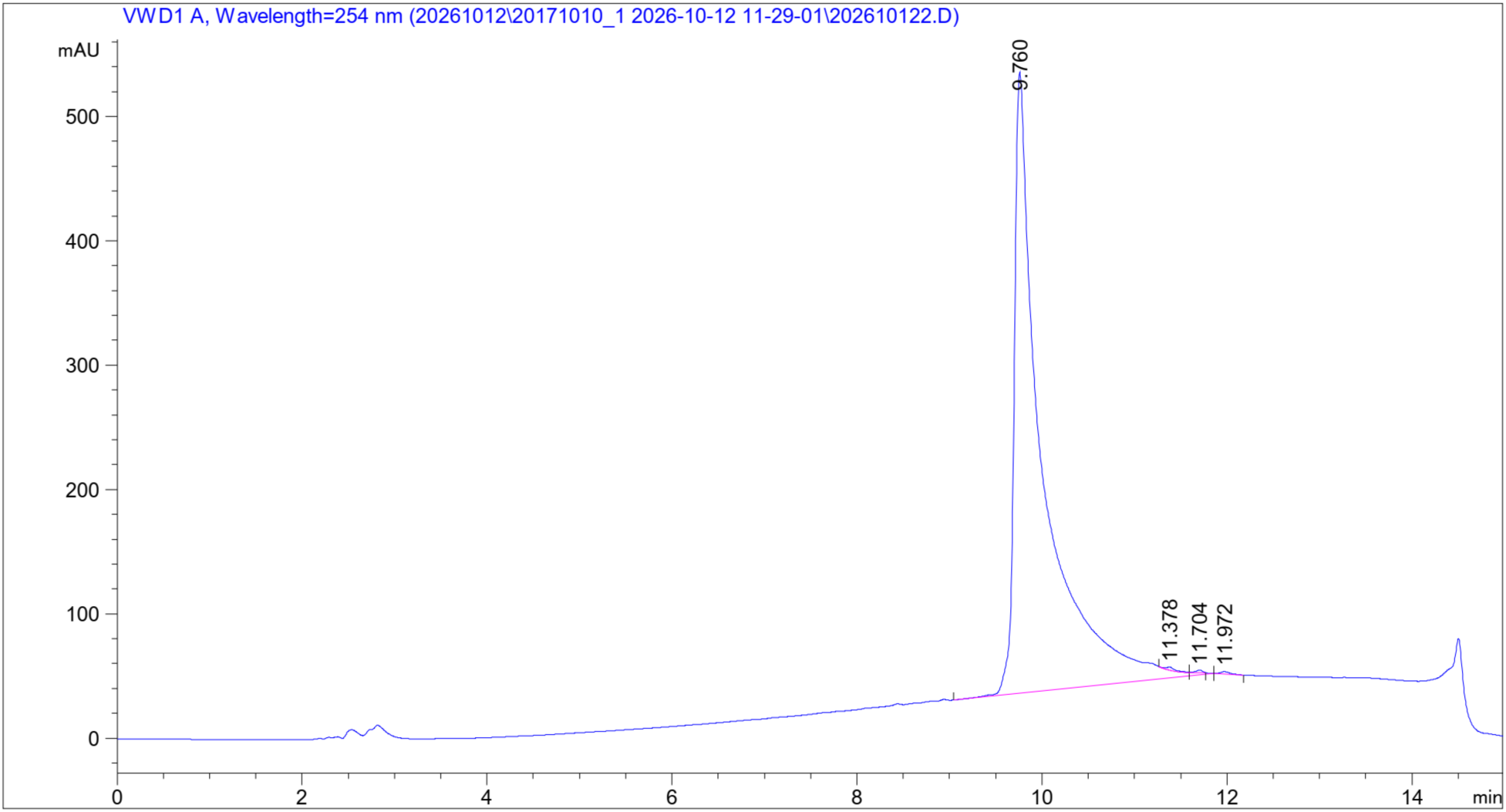
HPLC spectrum of Stearic acid-ATTO488.

**Figure S17.**
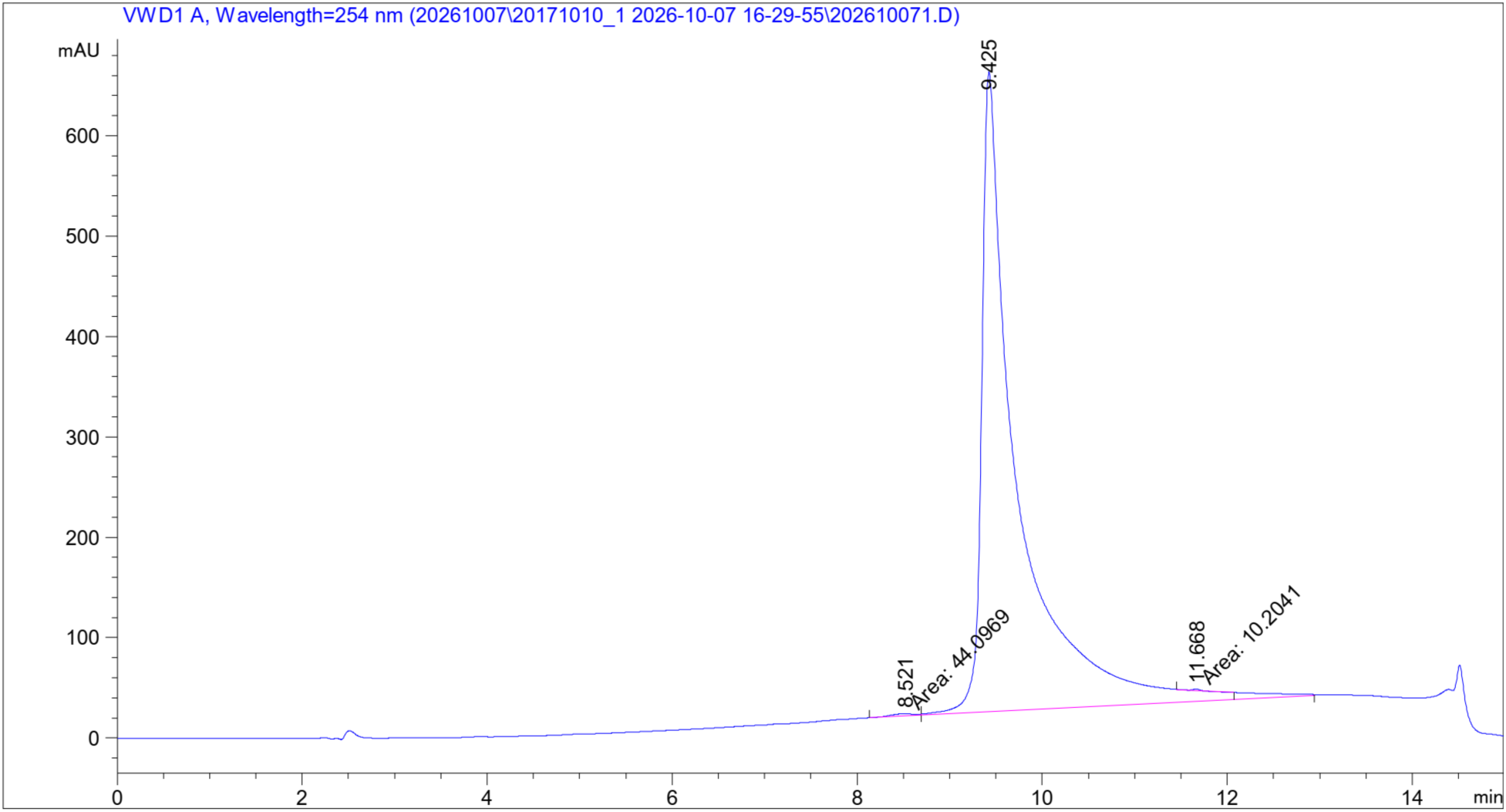
HPLC spectrum of BRD4-VHL-ATTO488.

## 2. Methods in Chemistry

### 2.1. General information

Chemicals that are commercially available were purchased from AmBeed, Combi-Block, Lumiprobe, MedChemExpress, TCI AMERICA, Sigma-Aldrich, and Thermo Fisher Scientific and used without further purification.

### 2.2. Nuclear magnetic resonance (NMR) spectroscopy

For NMR spectra, Proton NMR (^1^H-NMR) and carbon NMR (^13^C-NMR) were recorded on a Bruker Ascend Evo 400 spectrometer at ambient temperature. Chemical shifts (*δ*) are reported in ppm, coupling constants (*J*) in Hz, and signal multiplicities are denoted as follows: *s* (singlet), *d* (doublet), *t* (triplet), *q* (quartet), and *m* (multiplet).

### 2.3. High-performance liquid chromatography (HPLC)

Compound purity was determined by HPLC using an Agilent 1100 series system equipped with a Luna® Omega 5 μm Polar C_18_ column (100 Å, 150 × 4.6 mm). The column temperature was maintained at 37 °C. Elution was carried out at a flow rate of 0.8 mL/min using a mobile phase consisting of deionized water with 0.1% TFA (v/v) and methanol with 0.1% TFA (v/v).

### 2.4. Liquid chromatography–mass spectrometry (LC-MS) and ultra-high performance liquid chromatography-high resolution mass spectrometry (UHPLC–HRMS)

LC-MS analysis for reaction monitoring was performed on a Thermo Fisher LCQ-DECA mass spectrometer (ESI+ mode) equipped with a Luna® Omega 3 μm Polar C18 (100 Å, 50 × 3.0 mm) column at 35 °C. UHPLC–HRMS characterization of final compounds was conducted on an Orbitrap Exploris 120 mass spectrometer (ESI+ mode) using a Luna® Omega 5 μm Polar C18 (100 Å, 50 × 2.1 mm) column at 35 °C. For both LC–MS and UPLC–HRMS, elution was performed at a flow rate of 0.5 mL/min using a solvent system of deionized water containing 0.1% formic acid (v/v) and methanol containing 0.1% formic acid (v/v). For HPLC, LC–MS, and UHPLC–HRMS analyses, samples were dissolved in appropriate solvents (e.g., acetonitrile or methanol) and injected directly into the system using an automated sample handler.

### 2.5. Chemical synthesis

#### Step 1. Synthesis of 3,6-diamino-9-(2-((4-((17-carboxyheptadecyl)amino)-4-oxobutyl)(methyl)carbamoyl)phenyl)-4,5-disulfoxanthylium (2, Stearic acid-ATTO488)

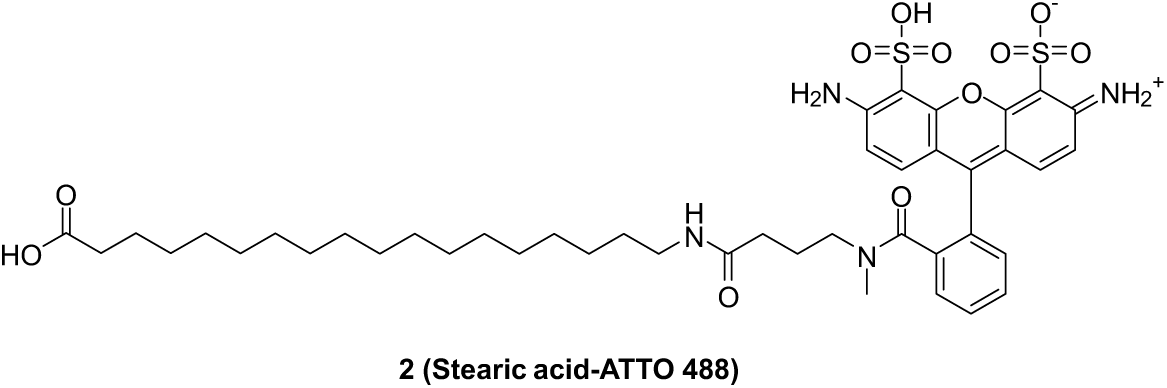

To a solution of 18-aminooctadecanoic acid (1, 2.18 mg, 0.00728 mmol, 1 equiv.) in dry dimethylformamide (1.5 mL) were added *N*,*N*-diisopropylethylamine (18.8 mg, 0.146 mmol, 20 equiv.) and ATTO488-NHS ester (5.0 mg, 0.00728 mmol, 1 equiv.). The reaction mixture was stirred at room temperature for 8 hours in darkness. The reaction mixture was diluted with methanol (3 mL) and purified via preparative HPLC on a C_18_ reverse-phase column (Luna® Omega 5 µm Polar C_18_ 100 Å, LC Column 250 x 4.6 mm, eluting with methanol/water (0.1% TFA in each, v/v)) and lyophilized to afford a red foam as 3,6-diamino-9-(2-((4-((17-carboxyheptadecyl)amino)-4-oxobutyl)(methyl)carbamoyl)phenyl)-4,5-disulfoxanthylium (**2**, **Stearic acid-ATTO488**, 2.4 mg, 37.8% yield).

**^1^H-NMR** (400 MHz, DMSO-*d_6_*) δ 9.07 (s, 2H), 8.69 – 8.35 (m, 2H), 7.76 – 7.32 (m, 5H), 7.05 – 6.87 (m, 4H), 3.50 (s, 3H), 3.05 – 3.01 (m, 2H), 2.92 – 2.86 (m, 2H), 2.77 – 2.70 (m, 2H), 2.05 – 1.89 (m, 3H), 1.72 – 1.58 (m, 2H), 1.39 – 1.14 (m, 30H).

**^13^C-NMR** (101 MHz, DMSO-*d_6_*) δ ^13^C NMR (101 MHz, DMSO) δ 171.75, 167.86, 158.59, 158.28, 157.28, 156.55, 155.81, 136.60, 132.00, 131.22, 130.66, 130.14, 129.88, 127.70, 119.33, 116.29, 113.74, 113.59, 49.07, 46.45, 37.58, 32.31, 29.48, 29.36, 29.20, 29.04, 26.86, 25.06, 22.45.

**HRMS** (ESI^+^) (*m/z*) calculated for C_43_H_58_N_4_O_11_S_2_, 870.3544; found 871.3599 [M+H]^+^.

**HPLC purity** 99.57%.

Step 2. Synthesis of (*S*)-2-(4-(4-chlorophenyl)-2,3,9-trimethyl-6*H*-thieno[3,2-*f*][1,2,4]triazolo[4,3-*a*][1,4]diazepin-6-yl)acetic acid (4, (+)-JQ1 acid)

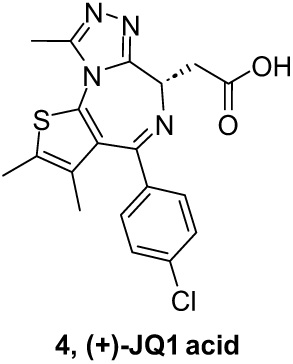

To a solution of (+)-JQ1 (**3**, 1.0 g, 2.19 mmol) in dichloromethane (15.0 mL) was added trifluoroacetic acid (5.0 mL). The reaction mixture was stirred at room temperature overnight. LC-MS analysis indicated the complete conversion. The reaction mixture was concentrated under vacuum, absorbed onto celite, and purified via C_18_ reversed flash chromatography (water: MeOH = 9: 1 to 100% MeOH) and concentrated under vacuum to afford a white solid as (*S*)-2-(4-(4-chlorophenyl)-2,3,9-trimethyl-6*H*-thieno[3,2-*f*][1,2,4]triazolo[4,3-*a*][1,4]diazepin-6-yl)acetic acid (**4**, **(+)-JQ1 acid**, 0.75 g, 85.3% yield).

**^1^H-NMR** (400 MHz, DMSO-*d_6_*) δ 7.49 (d, *J* = 8.4 Hz, 2H), 7.44 (d, *J* = 8.4 Hz, 2H), 4.46 (t, *J* = 7.0 Hz, 1H), 3.48 – 3.39 (m, 1H), 3.37 – 3.27 (m, 1H), 2.60 (s, 3H), 2.41 (s, 3H), 1.63 (s, 3H).

**HRMS** (ESI^+^) (*m/z*) calculated for C_19_H_17_ClN_4_O_2_S, 400.0761; found 401.0827 [M+H]^+^.

#### Step 3. Synthesis of 2-(2-(2-azidoethoxy)ethoxy)ethyl 4-methylbenzenesulfonate (6)

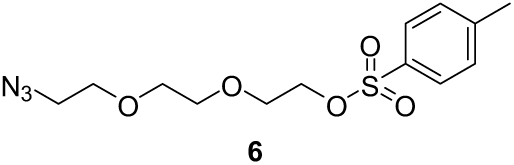

A mixture of triethylene glycol di(*p*-toluenesulfonate) (**5**, 25.0 g, 54.52 mmol, 1 equiv.) and sodium azide (3.5 g, 54.52 mmol, 1 equiv.) in ethanol (200.0 mL) was stirred at 80 ℃ overnight. After removing most of the ethanol under vacuum, ethyl acetate (100.0 mL) was added and resulting suspension was filtered through a short celite pad. The filtrate was concentrated under vacuum, absorbed onto silica gel, and purified via flash chromatography (hexane: ethyl acetate = 99: 1 to 1: 1) and concentrated under vacuum to afford a colorless oil as 2-(2-(2-azidoethoxy)ethoxy)ethyl 4-methylbenzenesulfonate (**6**, 5.42 g, 30.2% yield).

**^1^H-NMR** (400 MHz, DMSO-*d_6_*) δ 7.79 (d, *J* = 8.4 Hz, 2H), 7.48 (d, *J* = 8.5 Hz, 2H), 4.14 – 4.09 (m, 2H), 3.61 – 3.54 (m, 4H), 3.52 – 3.45 (m, 4H), 3.40 – 3.35 (m, 2H), 2.43 (s, 3H).

**HRMS** (ESI^+^) (*m/z*) calculated for C_13_H_19_N_3_O_5_S, 329.1045; found 330.1113 [M+H]^+^.

#### Step 4. Synthesis of 3-(2-(2-(2-azidoethoxy)ethoxy)ethoxy)-2-((2-(2-(2-azidoethoxy)ethoxy)ethoxy)methyl)-2-methylpropan-1-ol (7)

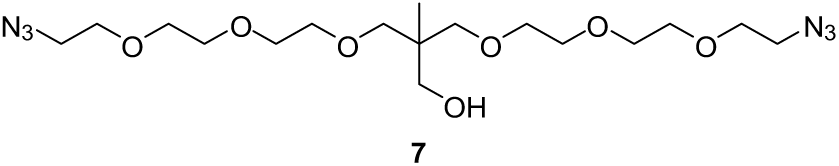

To a solution of trimethylolethane (0.99 g, 8.28 mmol, 1 equiv.) in dry dimethylformamide (20.0 mL) were added sodium hydride (in oil dispersion) 60% dispersion in mineral oil (0.73 g, 18.22 mmol, 2.2 equiv.) at 0 ℃. The reaction mixture was stirred at room temperature for 30 minutes. Then 2-(2-(2-azidoethoxy)ethoxy)ethyl 4-methylbenzenesulfonate (**6**, 5.42 g, 16.46 mmol, 2 equiv.) was added dropwise and the reaction mixture was stirred at 40 ℃ for 12 hours. After cooling to room temperature, the reaction mixture was diluted with ethyl acetate (200 mL) and ice-cold water (100 mL). The organic layer was washed with 0.1 N HCl aqueous solution and brine, dried over anhydrous sodium sulfate, filtered, concentrated under vacuum, absorbed onto silica gel, and purified via flash chromatography (hexane: ethyl acetate = 99: 1 to 100% ethyl acetate) and concentrated under vacuum to afford a colorless oil as 3-(2-(2-(2-azidoethoxy)ethoxy)ethoxy)-2-((2-(2-(2-azidoethoxy)ethoxy)ethoxy)methyl)-2-methylpropan-1-ol (**7**, 1.10 g, 30.6% yield).

**^1^H-NMR** (400 MHz, DMSO-*d_6_*) δ 4.30 (t, *J* = 5.3 Hz, 1H), 3.63 – 3.59 (m, 4H), 3.57 – 3.51 (m, 12H), 3.49 – 3.45 (m, 4H), 3.41 – 3.37 (m, 4H), 3.26 – 3.21 (m, 6H), 0.80 (s, 3H).

**HRMS** (ESI^+^) (*m/z*) calculated for C_17_H_34_N_6_O_7_, 434.2489; found 435.2557 [M+H]^+^.

#### Step 5. Synthesis of tert-butyl 1-azido-11-((2-(2-(2-azidoethoxy)ethoxy)ethoxy)methyl)-11-methyl-3,6,9,13-tetraoxapentadecan-15-oate (8)

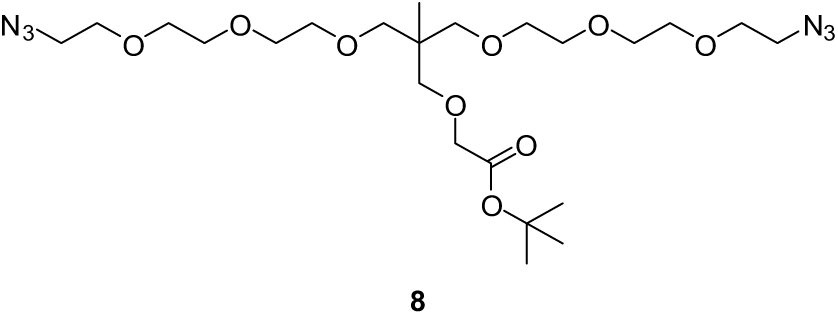

A mixture of 3-(2-(2-(2-azidoethoxy)ethoxy)ethoxy)-2-((2-(2-(2-azidoethoxy)ethoxy)ethoxy)methyl)-2-methylpropan-1-ol (**7**, 0.50 g, 1.15 mmol, 1 equiv.) and potassium *tert*-butoxide (0.32 g, 2.88 mmol, 2.5 equiv.) in dry tetrahydrofuran (10.0 mL) was stirred at 0 ℃ for 15 minutes. Then *tert*-butyl bromoacetate (0.34 g, 1.73 mmol, 1.5 equiv.) was added dropwise. The reaction mixture was stirred at room temperature overnight. The resulting mixture was concentrated under vacuum, absorbed onto silica gel, and purified via flash chromatography (hexane: ethyl acetate = 99: 1 to 100% ethyl acetate) and concentrated under vacuum to afford a light-yellow oil as *tert*-butyl 1-azido-11-((2-(2-(2-azidoethoxy)ethoxy)ethoxy)methyl)-11-methyl-3,6,9,13-tetraoxapentadecan-15-oate (**8**, 0.30 g, 47.6% yield).

**^1^H-NMR** (400 MHz, CDCl_3_) δ 3.86 (s, 2H), 3.63 – 3.54 (m, 16H), 3.50 (dd, *J* = 5.9, 3.8 Hz, 4H), 3.35 – 3.30 (m, 6H), 3.28 (s, 4H), 1.40 (s, 9H), 0.90 (s, 3H).

**HRMS** (ESI^+^) (*m/z*) calculated for C_23_H_44_N_6_O_9_, 548.3170; found 549.3220 [M+H]^+^.

#### Step 6. Synthesis of 1-azido-11-((2-(2-(2-azidoethoxy)ethoxy)ethoxy)methyl)-11-methyl-3,6,9,13-tetraoxapentadecan-15-oic acid (9)

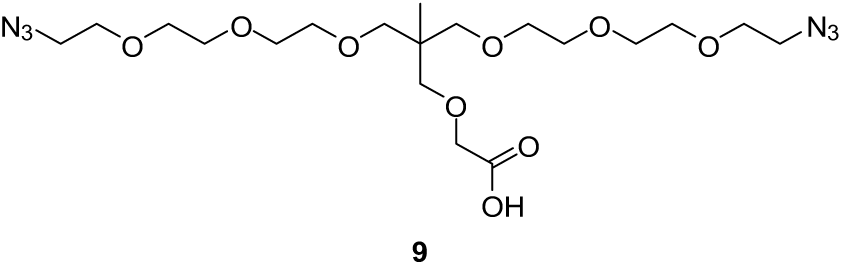

To a solution of *tert*-butyl 1-azido-11-((2-(2-(2-azidoethoxy)ethoxy)ethoxy)methyl)-11-methyl-3,6,9,13-tetraoxapentadecan-15-oate (**8**, 0.60 g, 1.09 mmol, 1 equiv.) in dichloromethane (12.0 mL) was added trifluoracetic acid (3.0 mL). The reaction mixture was stirred at room temperature overnight. LC-MS analysis indicated the completed conversion. The reaction mixture was concentrated under vacuum to afford a light-yellow oil as the crude product of 1-azido-11-((2-(2-(2-azidoethoxy)ethoxy)ethoxy)methyl)-11-methyl-3,6,9,13-tetraoxapentadecan-15-oic acid (**9**), which was used directly for the next steps without purifications.

**HRMS** (ESI^+^) (*m/z*) calculated for C_19_H_36_N_6_O_9_, 492.2544; found 493.2608 [M+H]^+^.

#### Step 7. Synthesis of (2S,4R)-1-((S)-1-azido-11-((2-(2-(2-azidoethoxy)ethoxy)ethoxy)methyl)-17-(tert-butyl)-11-methyl-15-oxo-3,6,9,13-tetraoxa-16-azaoctadecan-18-oyl)-4-hydroxy-N-((*S*)-1-(4-(4-methylthiazol-5-yl)phenyl)ethyl)pyrrolidine-2-carboxamide (10)

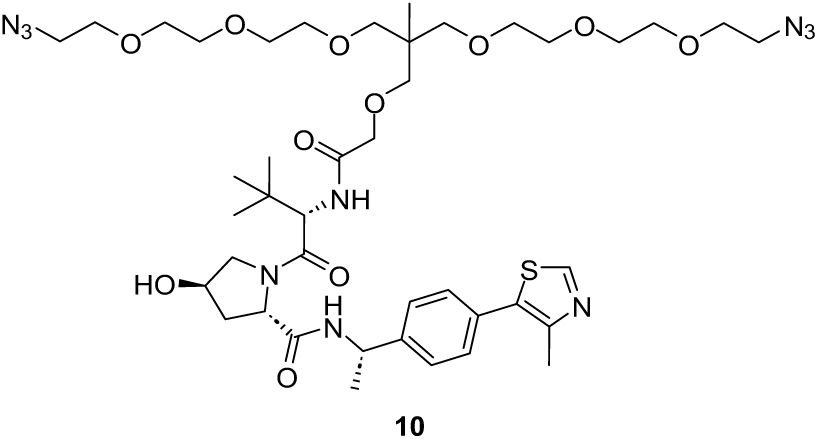

A mixture of 1-azido-11-((2-(2-(2-azidoethoxy)ethoxy)ethoxy)methyl)-11-methyl-3,6,9,13-tetraoxapentadecan-15-oic acid (**9**, 45.0 mg, 0.091 mmol, 1 equiv.), *N*,*N*-diisopropylethylamine (35.4 mg, 0.27 mmol, 3 equiv.), (*S*, *R*, *S*)-AHPC-Me hydrochloride (43.8 mg, 0.091 mmol, 1 equiv.), and 1-[bis(dimethylamino)methylene]-1*H*-1,2,3-triazolo[4,5-*b*]pyridinium 3-oxid hexafluorophosphate (34.7 mg, 0.091 mmol, 1 equiv.) in dichloromethane (1.0 mL) was stirred at room temperature overnight. Then the reaction mixture was concentrated under vacuum and purified via flash chromatography (dichloromethane: methanol = 99: 1 to 9: 1) and concentrated under vacuum to afford a colorless oil as (2*S*,4*R*)-1-((*S*)-1-azido-11-((2-(2-(2-azidoethoxy)ethoxy)ethoxy)methyl)-17-(*tert*-butyl)-11-methyl-15-oxo-3,6,9,13-tetraoxa-16-azaoctadecan-18-oyl)-4-hydroxy-*N*-((*S*)-1-(4-(4-methylthiazol-5-yl)phenyl)ethyl)pyrrolidine-2-carboxamide (**10**, 46.0 mg, 55.0% yield).

**^1^H-NMR** (400 MHz, DMSO-*d_6_*) δ 8.98 (s, 1H), 8.53 – 8.32 (m, 1H), 7.44 (d, *J* = 8.0 Hz, 2H), 7.37 (d, *J* = 7.9 Hz, 2H), 7.30 – 7.22 (m, 1H), 5.16 – 5.10 (m, 1H), 4.97 – 4.86 (m, 1H), 4.58 – 4.42 (m, 2H), 4.33 – 4.22 (m, 1H), 3.94 – 3.82 (m, 2H), 3.62 – 3.49 (m, 22H), 3.40 – 3.35 (m, 6H), 3.29 (d, *J* = 2.6 Hz, 4H), 2.46 (s, 3H), 2.11 – 2.00 (m, 1H), 1.84 – 1.70 (m, 1H), 1.38 (d, *J* = 6.7 Hz, 3H), 0.95 (s, 9H), 0.92 (s, 3H).

**HRMS** (ESI^+^) (*m/z*) calculated for C_42_H_66_N_10_O_11_S, 918.4633; found 919.4704 [M+H]^+^.

#### Step 8. Synthesis of (2S,4R)-1-((S)-1-amino-11-((2-(2-(2-aminoethoxy)ethoxy)ethoxy)methyl)-17-(tert-butyl)-11-methyl-15-oxo-3,6,9,13-tetraoxa-16-azaoctadecan-18-oyl)-4-hydroxy-N-((S)-1-(4-(4-methylthiazol-5-yl)phenyl)ethyl)pyrrolidine-2-carboxamide (11)

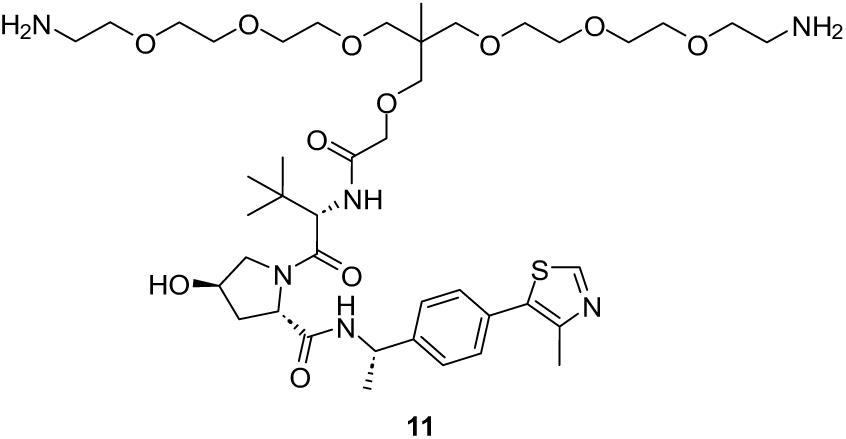

To a solution of (2*S*,4*R*)-1-((*S*)-1-azido-11-((2-(2-(2-azidoethoxy)ethoxy)ethoxy)methyl)-17-(*tert*-butyl)-11-methyl-15-oxo-3,6,9,13-tetraoxa-16-azaoctadecan-18-oyl)-4-hydroxy-*N*-((*S*)-1-(4-(4-methylthiazol-5-yl)phenyl)ethyl)pyrrolidine-2-carboxamide (**10**, 83.0 mg, 0.090 mmol, 1 equiv.) in methanol (1.0 mL) was added Pd/C (50.0 mg). The reaction mixture was stirred at room temperature overnight under hydrogen atmosphere. LC-MS analysis indicated the completed conversion. The reaction mixture was concentrated under vacuum to afford a light-yellow oil as the crude product of (2*S*,4*R*)-1-((*S*)-1-amino-11-((2-(2-(2-aminoethoxy)ethoxy)ethoxy)methyl)-17-(*tert*-butyl)-11-methyl-15-oxo-3,6,9,13-tetraoxa-16-azaoctadecan-18-oyl)-4-hydroxy-*N*-((*S*)-1-(4-(4-methylthiazol-5-yl)phenyl)ethyl)pyrrolidine-2-carboxamide (**11**), which was used directly for the next steps without purifications.

**HRMS** (ESI^+^) (*m/z*) calculated for C_42_H_70_N_6_O_11_S, 866.4823; found 867.4880 [M+H]^+^.

#### Step 9. Synthesis of (2S,4R)-1-((20S)-14-((2-(2-(2-aminoethoxy)ethoxy)ethoxy)methyl)-20-(tert-butyl)-1-((S)-4-(4-chlorophenyl)-2,3,9-trimethyl-6H-thieno[3,2-f][1,2,4]triazolo[4,3-a][1,4]diazepin-6-yl)-14-methyl-2,18-dioxo-6,9,12,16-tetraoxa-3,19-diazahenicosan-21-oyl)-4-hydroxy-N-((S)-1-(4-(4-methylthiazol-5-yl)phenyl)ethyl)pyrrolidine-2-carboxamide (12)

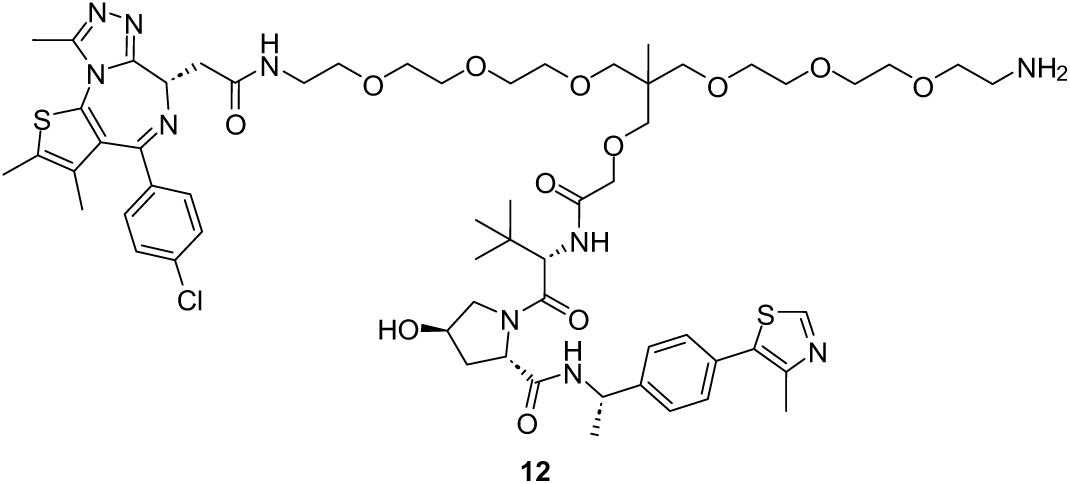

A solution of (2*S*,4*R*)-1-((*S*)-1-amino-11-((2-(2-(2-aminoethoxy)ethoxy)ethoxy)methyl)-17-(*tert*-butyl)-11-methyl-15-oxo-3,6,9,13-tetraoxa-16-azaoctadecan-18-oyl)-4-hydroxy-*N*-((*S*)-1-(4-(4-methylthiazol-5-yl)phenyl)ethyl)pyrrolidine-2-carboxamide (**11**, 78.0 mg, 0.090 mmol, 1 equiv.), *N*,*N*-diisopropylethylamine (77.6 mg, 0.40 mmol, 4.5 equiv.), (*S*)-2-(4-(4-chlorophenyl)-2,3,9-trimethyl-6*H*-thieno[3,2-*f*][1,2,4]triazolo[4,3-*a*][1,4]diazepin-6-yl)acetic acid (**4**, 34.3 mg, 0.086 mmol, 0.95 equiv.), 1-ethyl-3-(3-dimethylaminopropyl)carbodiimide hydrochloride (27.9 mg, 0.18 mmol, 2 equiv.), and 1-hydroxy-7-azabenzotriazole (24.5 mg, 0.18 mmol, 2 equiv.) in dry dimethylformamide (2.0 mL) was stirred at room temperature overnight. Then the reaction mixture was diluted with methanol (5.0 mL) and purified via preparative HPLC on a C_18_ reverse-phase column (Luna® Omega 5 µm Polar C_18_ 100 Å, LC Column 250 x 4.6 mm, eluting with methanol/water (0.1% TFA in each, v/v)) and lyophilized to afford a yellow foam as crude product of (2*S*,4*R*)-1-((20*S*)-14-((2-(2-(2-aminoethoxy)ethoxy)ethoxy)methyl)-20-(*tert*-butyl)-1-((*S*)-4-(4-chlorophenyl)-2,3,9-trimethyl-6*H*-thieno[3,2-*f*][1,2,4]triazolo[4,3-*a*][1,4]diazepin-6-yl)-14-methyl-2,18-dioxo-6,9,12,16-tetraoxa-3,19-diazahenicosan-21-oyl)-4-hydroxy-*N*-((*S*)-1-(4-(4-methylthiazol-5-yl)phenyl)ethyl)pyrrolidine-2-carboxamide (**12**, 25.1 mg, 22.3% yield), which was used directly for the next step without further purifications.

**HRMS** (ESI^+^) (*m/z*) calculated for C_61_H_85_ClN_10_O_12_S_2_, 1248.5478; found 1249.5531 [M+H]^+^.

#### Step 10. Synthesis of 3,6-diamino-9-(2-((1-((S)-4-(4-chlorophenyl)-2,3,9-trimethyl-6H-thieno[3,2-f][1,2,4]triazolo[4,3-a][1,4]diazepin-6-yl)-14-((2-(((S)-1-((2S,4R)-4-hydroxy-2-(((S)-1-(4-(4-methylthiazol-5-yl)phenyl)ethyl)carbamoyl)pyrrolidin-1-yl)-3,3-dimethyl-1-oxobutan-2-yl)amino)-2-oxoethoxy)methyl)-14-methyl-2,26-dioxo-6,9,12,16,19,22-hexaoxa-3,25-diazanonacosan-29-yl)(methyl)carbamoyl)phenyl)-4,5-disulfoxanthylium (13, BRD4-VHL-ATTO488)

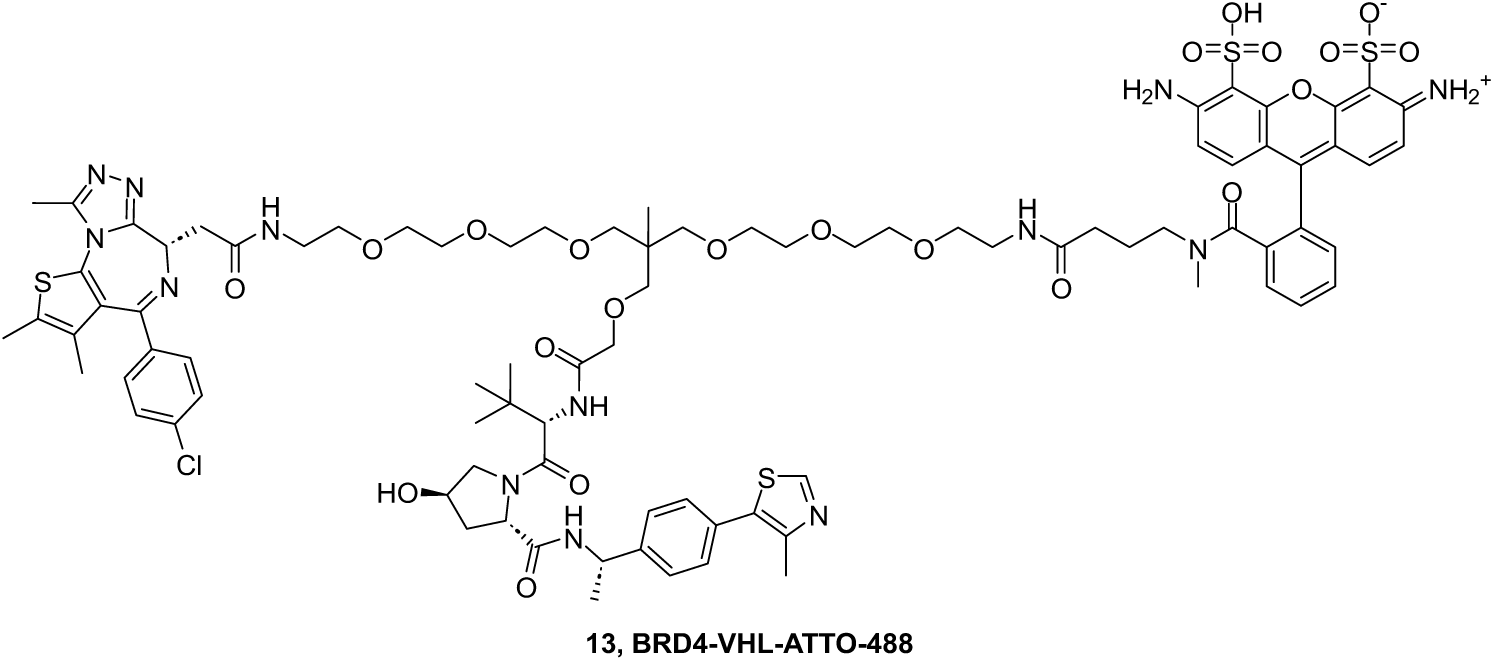

To a solution of (2*S*,4*R*)-1-((20*S*)-14-((2-(2-(2-aminoethoxy)ethoxy)ethoxy)methyl)-20-(*tert*-butyl)-1-((*S*)-4-(4-chlorophenyl)-2,3,9-trimethyl-6*H*-thieno[3,2-*f*][1,2,4]triazolo[4,3-*a*][1,4]diazepin-6-yl)-14-methyl-2,18-dioxo-6,9,12,16-tetraoxa-3,19-diazahenicosan-21-oyl)-4-hydroxy-*N*-((*S*)-1-(4-(4-methylthiazol-5-yl)phenyl)ethyl)pyrrolidine-2-carboxamide (**12**, 36.4 mg, 0.029 mmol, 1 equiv.) in dry dimethylformamide (1.0 mL) were added *N*,*N*-diisopropylethylamine (75.0 mg, 0.58 mmol, 20 equiv.) and ATTO488-NHS ester (20.0 mg, 0.029 mmol, 1 equiv.) The reaction mixture was stirred at room temperature for 4 hours in darkness. LC-MS analysis indicated the completed conversion of the reaction. The reaction mixture was diluted with water (5 mL) and purified via preparative HPLC on a C_18_ reverse-phase column (Luna® Omega 5 µm Polar C_18_ 100 Å, LC Column 250 x 4.6 mm, eluting with methanol/water (0.1% TFA in each, v/v)) and lyophilized to afford a red foam as 3,6-diamino-9-(2-((1-((*S*)-4-(4-chlorophenyl)-2,3,9-trimethyl-6*H*-thieno[3,2-*f*][1,2,4]triazolo[4,3-*a*][1,4]diazepin-6-yl)-14-((2-(((*S*)-1-((2*S*,4*R*)-4-hydroxy-2-(((*S*)-1-(4-(4-methylthiazol-5-yl)phenyl)ethyl)carbamoyl)pyrrolidin-1-yl)-3,3-dimethyl-1-oxobutan-2-yl)amino)-2-oxoethoxy)methyl)-14-methyl-2,26-dioxo-6,9,12,16,19,22-hexaoxa-3,25-diazanonacosan-29-yl)(methyl)carbamoyl)phenyl)-4,5-disulfoxanthylium (**13**, **BRD4-VHL-ATTO488**, 13.9 mg, 26.3% yield).

**^1^H-NMR** (400 MHz, DMSO-*d_6_*) δ 9.03 (s, 2H), 8.92 (s, 1H), 8.72 – 8.30 (m, 3H), 8.21 (t, *J* = 5.6 Hz, 1H), 7.83 – 7.46 (m, 4H), 7.50 – 7.30 (m, 6H), 7.29 (d, *J* = 7.8 Hz, 2H), 7.19 (d, *J* = 9.4 Hz, 1H), 7.00 (d, *J* = 9.2 Hz, 2H), 6.92 (d, *J* = 9.3 Hz, 2H), 4.89 – 4.77 (m, 1H), 4.57 – 4.27 (m, 3H), 4.24 – 4.15 (m, 1H), 3.97 – 2.96 (m, 40H), 2.75 (s, 2H), 2.62 – 2.54 (m, 1H), 2.52 (s, 3H), 2.38 (s, 3H), 2.33 (s, 3H), 2.06 – 1.89 (m, 2H), 1.79 – 1.57 (m, 3H), 1.54 (s, 3H), 1.33 (s, 1H), 1.30 (d, *J* = 7.0 Hz, 3H), 1.17 (s, 1H), 0.87 (s, 9H), 0.83 (s, 3H).

**^13^C-NMR** (101 MHz, DMSO-*d_6_*) δ ^13^C NMR (101 MHz, DMSO) δ 171.92, 170.90, 170.13, 169.48, 168.77, 167.84, 163.50, 157.28, 156.58, 155.67, 155.58, 151.91, 150.29, 148.19, 145.23, 137.22, 136.67, 135.72, 132.71, 132.01, 131.60, 131.23, 130.62, 130.32, 130.15, 130.08, 129.80, 129.35, 129.29, 128.92, 127.65, 126.78, 119.34, 113.62, 74.25, 73.52, 70.99, 70.70, 70.31, 70.26, 70.12, 70.01, 69.70, 69.51, 69.23, 58.98, 57.01, 56.07, 54.30, 48.28, 38.23, 37.51, 36.34, 32.23, 26.67, 22.89, 22.44, 17.70, 16.43, 14.49, 13.14, 11.74.

**HRMS** (ESI^+^) (*m/z*) calculated for C_86_H_106_ClN_13_O_21_S_4_, 1819.6198; found 910.8174 [M+2H]^2+^.

**HPLC purity** 99.69%.

## References

1. Pepino, M. Y., Kuda, O., Samovski, D. & Abumrad, N. A. Structure-Function of CD36 and Importance of Fatty Acid Signal Transduction in Fat Metabolism. Annu. Rev. Nutr. 34, 281–303 (2014).

2. Silverstein, R. L. & Febbraio, M. CD36, a Scavenger Receptor Involved in Immunity, Metabolism, Angiogenesis, and Behavior. Sci. Signal. 2, (2009).

3. Greenwalt, D. et al. Membrane glycoprotein CD36: a review of its roles in adherence, signal transduction, and transfusion medicine. Blood 80, 1105–1115 (1992).

4. Zeng, Y., Tao, N., Chung, K.-N., Heuser, J. E. & Lublin, D. M. Endocytosis of Oxidized Low Density Lipoprotein through Scavenger Receptor CD36 Utilizes a Lipid Raft Pathway That Does Not Require Caveolin-1. J. Biol. Chem. 278, 45931–45936 (2003).

5. Hames, K. C., Vella, A., Kemp, B. J. & Jensen, M. D. Free Fatty Acid Uptake in Humans With CD36 Deficiency. Diabetes 63, 3606–3614 (2014).

6. Kennedy, D. J. et al. A CD36-dependent pathway enhances macrophage and adipose tissue inflammation and impairs insulin signalling. Cardiovasc. Res. 89, 604–613 (2011).

7. Aitman, T. J. et al. Identification of Cd36 (Fat) as an insulin-resistance gene causing defective fatty acid and glucose metabolism in hypertensive rats. Nat. Genet. 21, 76–83 (1999).

8. Susztak, K., Ciccone, E., McCue, P., Sharma, K. & Böttinger, E. P. Multiple Metabolic Hits Converge on CD36 as Novel Mediator of Tubular Epithelial Apoptosis in Diabetic Nephropathy. PLoS Med. 2, e45 (2005).

9. Wang, J. & Li, Y. CD36 tango in cancer: signaling pathways and functions. Theranostics 9, 4893–4908 (2019).

10. Guerrero-Rodríguez, S. L., Mata-Cruz, C., Pérez-Tapia, S. M. & Velasco-Velázquez, M. A. Role of CD36 in cancer progression, stemness, and targeting. Front. Cell Dev. Biol. 10, 1079076 (2022).

11. Kim, C. H. Short-, medium-, versus long-chain fatty acids: mechanisms of immunomodulation and disease pathogenesis. Cell. Mol. Immunol. 23, 594–606 (2026).

12. Zhang, S., et al. Long-chain fatty acyl coenzyme A inhibits NME1/2 and regulates cancer metastasis. Proc. Natl. Acad. Sci. 119, e2117013119 (2022).

13. Akiba, Y., Nishii, S., Mizoguchi, A., Ito, S. & Kaunitz, J. D. Lipopolysaccharide transport during long-chain fatty acid exposure is mediated by caveolin-1 dependent endocytosis in murine jejunum. Tissue Barriers 13, 2482277 (2025).

14. Wang, Z. et al. CD36-mediated endocytosis of proteolysis-targeting chimeras. Cell 188, 3219–3237. e18 (2025).

15. Burslem, G. M. & Crews, C. M. Proteolysis-targeting chimeras as therapeutics and tools for biological discovery. Cell 181, 102–114 (2020).

16. Békés, M., Langley, D. R. & Crews, C. M. PROTAC targeted protein degraders: the past is prologue. Nat. Rev. Drug Discov. 21, 181–200 (2022).

17. Donovan, K. A. et al. Mapping the degradable kinome provides a resource for expedited degrader development. Cell 183, 1714–1731 (2020).

18. Han, X. & Sun, Y. Strategies for the discovery of oral PROTAC degraders aimed at cancer therapy. Cell Rep. Phys. Sci. 3, (2022).

19. Hao, J.-W. et al. CD36 facilitates fatty acid uptake by dynamic palmitoylation-regulated endocytosis. Nat. Commun. 11, 4765 (2020).

20. Ring, A., Le Lay, S., Pohl, J., Verkade, P. & Stremmel, W. Caveolin-1 is required for fatty acid translocase (FAT/CD36) localization and function at the plasma membrane of mouse embryonic fibroblasts. Biochim. Biophys. Acta BBA - Mol. Cell Biol. Lipids 1761, 416–423 (2006).

21. Pohl, J., Ring, A., Korkmaz, Ü., Ehehalt, R. & Stremmel, W. FAT/CD36-mediated Long-Chain Fatty Acid Uptake in Adipocytes Requires Plasma Membrane Rafts. Mol. Biol. Cell 16, 24–31 (2005).

22. Stoeber, M. et al. Oligomers of the ATPase EHD2 confine caveolae to the plasma membrane through association with actin. EMBO J. 31, 2350–2364 (2012).

23. Thomsen, P., Roepstorff, K., Stahlhut, M. & Van Deurs, B. Caveolae Are Highly Immobile Plasma Membrane Microdomains, Which Are not Involved in Constitutive Endocytic Trafficking. Mol. Biol. Cell 13, 238–250 (2002).

24. Bitsikas, V., Corrêa, I. R. & Nichols, B. J. Clathrin-independent pathways do not contribute significantly to endocytic flux. eLife 3, e03970 (2014).

25. Shvets, E., Bitsikas, V., Howard, G., Hansen, C. G. & Nichols, B. J. Dynamic caveolae exclude bulk membrane proteins and are required for sorting of excess glycosphingolipids. Nat. Commun. 6, 6867 (2015).

26. Hamilton, J. A., Guo, W. & Kamp, F. Mechanism of cellular uptake of long-chain fatty acids: Do we need cellular proteins? in Cellular Lipid Binding Proteins (ed. Glatz, J. F. C.) 17–23 (Springer US, Boston, MA, 2002). doi:10.1007/978-1-4419-9270-3_3.

27. Doege, H. & Stahl, A. Protein-Mediated Fatty Acid Uptake: Novel Insights from In Vivo Models. Physiology 21, 259–268 (2006).

28. Black, P. N. & DiRusso, C. C. Transmembrane Movement of Exogenous Long-Chain Fatty Acids: Proteins, Enzymes, and Vectorial Esterification. Microbiol. Mol. Biol. Rev. 67, 454–472 (2003).

29. Stahl, A. A current review of fatty acid transport proteins (SLC27). Pfl◻gers Arch. Eur. J. Physiol. 447, 722–727 (2004).

30. Bonen, A. et al. Plasmalemmal fatty acid transport is regulated in heart and skeletal muscle by contraction, insulin and leptin, and in obesity and diabetes. Acta Physiol. Scand. 178, 347–356 (2003).

31. Schaffer, J. E. & Lodish, H. F. Expression cloning and characterization of a novel adipocyte long chain fatty acid transport protein. Cell 79, 427–436 (1994).

32. Wu, Q. et al. FATP1 Is an Insulin-Sensitive Fatty Acid Transporter Involved in Diet-Induced Obesity. Mol. Cell. Biol. 26, 3455–3467 (2006).

33. Matsson, P. & Kihlberg, J. How Big Is Too Big for Cell Permeability? Journal of medicinal chemistry vol. 60 1662–1664 (ACS Publications, 2017).

34. Scott, D. E. et al. Systematic investigation of the permeability of androgen receptor PROTACs. ACS Med. Chem. Lett. 11, 1539–1547 (2020).

35. Lipinski, C. A., Lombardo, F., Dominy, B. W. & Feeney, P. J. Experimental and computational approaches to estimate solubility and permeability in drug discovery and development settings. Adv. Drug Deliv. Rev. 23, 3–25 (1997).

36. Hayer, A., Stoeber, M., Bissig, C. & Helenius, A. Biogenesis of Caveolae: Stepwise Assembly of Large Caveolin and Cavin Complexes. Traffic 11, 361–382 (2010).

37. Tagawa, A. et al. Assembly and trafficking of caveolar domains in the cell. J. Cell Biol. 170, 769–779 (2005).

38. Traub, L. M. Tickets to ride: selecting cargo for clathrin-regulated internalization. Nat. Rev. Mol. Cell Biol. 10, 583–596 (2009).

39. Aguet, F. et al. Membrane dynamics of dividing cells imaged by lattice light-sheet microscopy. Mol. Biol. Cell 27, 3418–3435 (2016).

40. Blondeau, F. et al. Tandem MS analysis of brain clathrin-coated vesicles reveals their critical involvement in synaptic vesicle recycling. Proc. Natl. Acad. Sci. 101, 3833–3838 (2004).

41. Grimm, J. B. et al. A general method to improve fluorophores for live-cell and single-molecule microscopy. Nat. Methods 12, 244–250 (2015).

42. Harding, C., Heuser, J. & Stahl, P. Receptor-mediated endocytosis of transferrin and recycling of the transferrin receptor in rat reticulocytes. J. Cell Biol. 97, 329–339 (1983).

43. Collawn, J. F. et al. Transferrin receptor internalization sequence YXRF implicates a tight turn as the structural recognition motif for endocytosis. Cell 63, 1061–1072 (1990).

44. DeGroot, A. C. M., Gollapudi, S., Zhao, C., LaMonica, M. F. & Stachowiak, J. C. Weakly Internalized Receptors Use Coated Vesicle Heterogeneity to Evade Competition during Endocytosis. Biochemistry 60, 2195–2205 (2021).

45. McGraw, T. E., Pytowski, B., Arzt, J. & Ferrone, C. Mutagenesis of the human transferrin receptor: two cytoplasmic phenylalanines are required for efficient internalization and a second-site mutation is capable of reverting an internalization-defective phenotype. J. Cell Biol. 112, 853–861 (1991).

46. Zhao, C. et al. Receptor heterodimerization modulates endocytosis through collaborative and competitive mechanisms. Biophys. J. 117, 646–658 (2019).

47. Aguet, F., Antonescu, C. N., Mettlen, M., Schmid, S. L. & Danuser, G. Advances in Analysis of Low Signal-to-Noise Images Link Dynamin and AP2 to the Functions of an Endocytic Checkpoint. Dev. Cell 26, 279–291 (2013).

48. DeGroot, A. C. M. et al. Entropic Control of Receptor Recycling Using Engineered Ligands. Biophys. J. 114, 1377–1388 (2018).

49. Shu, H. et al. The role of CD36 in cardiovascular disease. Cardiovasc. Res. 118, 115–129 (2022).

50. Martin, C. et al. CD36 as a lipid sensor. Physiol. Behav. 105, 36–42 (2011).

51. Pepino, M. Y., Kuda, O., Samovski, D. & Abumrad, N. A. Structure-Function of CD36 and Importance of Fatty Acid Signal Transduction in Fat Metabolism. Annu. Rev. Nutr. 34, 281–303 (2014).

52. Dores, M. R., Schnell, J. D., Maldonado-Baez, L., Wendland, B. & Hicke, L. The Function of Yeast Epsin and Ede1 Ubiquitin-Binding Domains During Receptor Internalization. Traffic 11, 151–160 (2010).

53. Klapisz, E. et al. A Ubiquitin-interacting Motif (UIM) Is Essential for Eps15 and Eps15R Ubiquitination. J. Biol. Chem. 277, 30746–30753 (2002).

54. Smith, J., Su, X., El-Maghrabi, R., Stahl, P. D. & Abumrad, N. A. Opposite Regulation of CD36 Ubiquitination by Fatty Acids and Insulin. J. Biol. Chem. 283, 13578–13585 (2008).

55. A Structural Explanation for the Recognition of Tyrosine-Based Endocytotic Signals. https://www.science.org/doi/10.1126/science.282.5392.1327 doi:10.1126/science.282.5392.1327.

56. Géminard, C., de Gassart, A., Blanc, L. & Vidal, M. Degradation of AP2 During Reticulocyte Maturation Enhances Binding of Hsc70 and Alix to a Common Site on TfR for Sorting into Exosomes. Traffic 5, 181–193 (2004).

57. Takai, M. et al. Unsaturated long-chain fatty acids inhibit the binding of oxidized low-density lipoproteins to a model CD36. Biosci. Biotechnol. Biochem. 78, 238–244 (2014).

58. Kuda, O. et al. Sulfo-N-succinimidyl Oleate (SSO) Inhibits Fatty Acid Uptake and Signaling for Intracellular Calcium via Binding CD36 Lysine 164. J. Biol. Chem. 288, 15547–15555 (2013).

59. Hughes, L. D., Rawle, R. J. & Boxer, S. G. Choose Your Label Wisely: Water-Soluble Fluorophores Often Interact with Lipid Bilayers. PLoS ONE 9, e87649 (2014).

60. Hopkins, C. R. & Trowbridge, I. S. Internalization and processing of transferrin and the transferrin receptor in human carcinoma A431 cells. J. Cell Biol. 97, 508–521 (1983).

61. Dejonghe, W. et al. Disruption of endocytosis through chemical inhibition of clathrin heavy chain function. Nat. Chem. Biol. 15, 641–649 (2019).

62. Goldstein, J. L., Basu, S. K., Brunschede, G. Y. & Brown, M. S. Release of low density lipoprotein from its cell surface receptor by sulfated glycosaminoglycans. Cell 7, 85–95 (1976).

63. Kamp, F., Zakim, D., Zhang, F., Noy, N. & Hamilton, J. A. Fatty Acid Flip-Flop in Phospholipid Bilayers Is Extremely Fast. Biochemistry 34, 11928–11937 (1995).

64. Klein, V. G. et al. Understanding and improving the membrane permeability of VH032-based PROTACs. ACS Med. Chem. Lett. 11, 1732–1738 (2020).

65. Gadd, M. S. et al. Structural basis of PROTAC cooperative recognition for selective protein degradation. Nat. Chem. Biol. 13, 514–521 (2017).

66. Zhao, X. et al. Expression of auxilin or AP180 inhibits endocytosis by mislocalizing clathrin: evidence for formation of nascent pits containing AP1 or AP2 but not clathrin. J. Cell Sci. 114, 353–365 (2001).

67. Neklesa, T. et al. ARV-110: An oral androgen receptor PROTAC degrader for prostate cancer. J. Clin. Oncol. 37, 259–259 (2019).

68. Silverstein, R. L. & Febbraio, M. CD36, a Scavenger Receptor Involved in Immunity, Metabolism, Angiogenesis, and Behavior. Sci. Signal. 2, (2009).

69. Polo, S. et al. A single motif responsible for ubiquitin recognition and monoubiquitination in endocytic proteins. Nature 416, 451–455 (2002).

70. Piper, R. C., Dikic, I. & Lukacs, G. L. Ubiquitin-dependent sorting in endocytosis. Cold Spring Harb. Perspect. Biol. 6, a016808 (2014).

71. Shih, S. C. et al. Epsins and Vps27p/Hrs contain ubiquitin-binding domains that function in receptor endocytosis. Nat. Cell Biol. 4, 389–393 (2002).

72. Kazazic, M. et al. Epsin 1 is Involved in Recruitment of Ubiquitinated EGF Receptors into Clathrin-Coated Pits. Traffic 10, 235–245 (2009).

73. Hawryluk, M. J. et al. Epsin 1 is a polyubiquitin-selective clathrin-associated sorting protein. Traffic 7, 262–281 (2006).

74. Boll, W. et al. Sequence requirements for the recognition of tyrosine-based endocytic signals by clathrin AP-2 complexes. EMBO J. 15, 5789–5795 (1996).

75. Xu, S., Jay, A., Brunaldi, K., Huang, N. & Hamilton, J. A. CD36 Enhances Fatty Acid Uptake by Increasing the Rate of Intracellular Esterification but Not Transport across the Plasma Membrane. Biochemistry 52, 7254–7261 (2013).

76. Son, N.-H. et al. Endothelial cell CD36 optimizes tissue fatty acid uptake. J. Clin. Invest. 128, 4329–4342 (2018).

77. Glatz, J. F. C., Luiken, J. J. F. P., Van Nieuwenhoven, F. A. & Van Der Vusse, G. J. Molecular mechanism of cellular uptake and intracellular translocation of fatty acids. Prostaglandins Leukot. Essent. Fatty Acids 57, 3–9 (1997).

78. Schmid, S. L. Reciprocal regulation of signaling and endocytosis: Implications for the evolving cancer cell. J. Cell Biol. 216, 2623–2632 (2017).

79. Mellman, I. & Yarden, Y. Endocytosis and Cancer. Cold Spring Harb. Perspect. Biol. 5, a016949–a016949 (2013).

80. Reis, C. R., Chen, P.-H., Bendris, N. & Schmid, S. L. TRAIL-death receptor endocytosis and apoptosis are selectively regulated by dynamin-1 activation. Proc. Natl. Acad. Sci. 114, 504–509 (2017).

81. Collawn, J. F. et al. Transferrin receptor internalization sequence YXRF implicates a tight turn as the structural recognition motif for endocytosis. Cell 63, 1061–1072 (1990).

82. Sorkin, A. & Goh, L. K. Endocytosis and intracellular trafficking of ErbBs. Exp. Cell Res. 315, 683–696 (2009).

83. Sorkina, T., Huang, F., Beguinot, L. & Sorkin, A. Effect of Tyrosine Kinase Inhibitors on Clathrin-coated Pit Recruitment and Internalization of Epidermal Growth Factor Receptor. J. Biol. Chem. 277, 27433–27441 (2002).

84. Wolfe, B. L. & Trejo, J. Clathrin-Dependent Mechanisms of G Protein-coupled Receptor Endocytosis. Traffic 8, 462–470 (2007).

85. Ferreira, F. et al. Endocytosis of G Protein-Coupled Receptors Is Regulated by Clathrin Light Chain Phosphorylation. Curr. Biol. 22, 1361–1370 (2012).

86. Zhang, Y. & Allison, J. P. Interaction of CTLA-4 with AP50, a clathrin-coated pit adaptor protein. Proc. Natl. Acad. Sci. 94, 9273–9278 (1997).

87. Qureshi, O. S. et al. Constitutive Clathrin-mediated Endocytosis of CTLA-4 Persists during T Cell Activation. J. Biol. Chem. 287, 9429–9440 (2012).

88. McKinnon, K. M. Flow Cytometry: An Overview. Curr. Protoc. Immunol. 120, (2018).

89. Gollapudi, S., et al. Steric pressure between glycosylated transmembrane proteins inhibits internalization by endocytosis. Proc. Natl. Acad. Sci. 120, e2215815120 (2023).

